# Targeting CD38 effectively prevents myelofibrosis in myeloproliferative neoplasms

**DOI:** 10.1101/2025.03.02.639410

**Authors:** Yiru Yan, Jinqin Liu, Songyang Zhao, Fuhui Li, Lin Yang, Zefeng Xu, Tiejun Qin, Xiaofan Zhu, Wenbin An, Gang Huang, Raajit K. Rampal, Zhijian Xiao, Bing Li

**Affiliations:** State Key Laboratory of Experimental Hematology, National Clinical Research Center for Blood Diseases, Haihe Laboratory of Cell Ecosystem, Institute of Hematology & Blood Diseases Hospital, Chinese Academy of Medical Sciences & Peking Union Medical College, Tianjin 300020, China; Tianjin Institutes of Health Science, Tianjin 301600, China; MDS and MPN Centre, Institute of Hematology and Blood Diseases Hospital, Chinese Academy of Medical Sciences & Peking Union Medical College, Tianjin, China; Department of pediatric hematology and oncology, Institute of Hematology and Blood Diseases Hospital, Chinese Academy of Medical Sciences & Peking Union Medical College, Tianjin, China; Department of Cell Systems and Anatomy, Department of Pathology and Laboratory Medicine, UT Health San Antonio, Joe R. and Teresa Lozano Long School of Medicine, Mays Cancer Center at UT Health San Antonio, San Antonio, Texas, USA; Department of Medicine, Memorial Sloan Kettering Cancer Center, New York, USA; Hematologic Pathology Center, Institute of Hematology and Blood Diseases Hospital, Chinese Academy of Medical Sciences & Peking Union Medical College, Tianjin, China

## Abstract

Proinflammatory signaling is a hallmark of myeloproliferative neoplasms (MPNs). Several studies have shown that monocytes are a major source of proinflammatory cytokines and monocyte-derived fibrocytes play a pivotal role in the pathogenesis of myelofibrosis (MF). To further explore the role of monocytes in MF, we generated inducible *Nras^G12D/+^Jak2^V617F/+^*(NJ) mice. Recipients transplanted with NJ bone marrow cells developed MF with an early onset of anemia and monocytosis. *In vitro*, NJ recipients’ bone marrow nucleated cells exhibited increased quantity of CD45^+^CollagenI^+^ fibrocytes, which were mainly derived from the Ly6c^high^ monocytes. RNA sequencing identified a significant elevated expression of CD38 (a nicotinamide adenine dinucleotide (NAD)^+^ hydrolase) in Ly6c^high^ monocytes from NJ mice, which results in pronounced lower level of NAD^+^. In humans, CD14^+^ monocytes from MF patients showed significantly higher expression of CD38 than controls and monocytes from polycythemia vera (PV) patients with grade 1 fibrosis had higher CD38 expression than those without fibrosis. Finally, we tested that boosting NAD^+^ via pharmacological CD38 targeting or NAD^+^ precursor supplementation inhibited the differentiation of fibrocytes *in vitro* and observed that targeting CD38 can effectively prevent the onset of fibrosis *in vivo*. Collectively, our findings shed light on the role of CD38 in monocytes and suggest potential clinical applications such as use of CD38 as a biomarker of fibrotic progression and potential clinical utility of CD38 inhibition in patients with MF.

**Key Points:**

1. CD38-overexpressing monocytes are increased in MF murine models and MPN patients progressing to fibrotic-phase disease.
2. Restoring intracellular NAD^+^ levels using the CD38 inhibitor 78c prevented the development of fibrotic-phase disease in MPN murine models.

## Introduction

BCR-ABL-negative myeloproliferative neoplasms (MPN) include polycythemia vera (PV), essential thrombocytosis (ET) and primary myelofibrosis (PMF). These disease are characterized by activating mutations in the JAK-STAT pathway, most commonly *JAK2*^V617F^.^1–4^ PMF, which carries the worst prognosis of these diseases, is characterized by bone marrow (BM) reticulin/collagen fibrosis, aberrant inflammatory cytokine expression, anemia, and extramedullary hematopoiesis (EMH).^5^ Approximately 15% of patients with ET or PV might progress into post-ET/PV myelofibrosis (MF), with similar outcome to PMF.^6, 7^ Importantly, BM fibrosis grade ≥ 2 (on a scale of 0-3)^8^ has been identified as an independent predictor of survival and is included in the Mutation-Enhanced International Prognostic Score System (MIPSS70) as well as karyotype-enhanced MIPSS70 (MIPSS70+).^9, 10^ While JAK inhibitors have some therapeutic benefits, such as decrease of splenomegaly and symptom alleviation, its efficacy in reversing fibrosis is limited.^11–13^ Therefore, it is essential to identify alternative mechanisms and targets of fibrotic progression for and to identify new biomarkers that can predict progression to secondary MF.

Inflammation has been identified as a crucial factor in the onset and advancement of MF.^14–18^ Aberrant immature megakaryocytes and platelet^19–23^, as well as monocytes, are the major origin of elevated cytokines in MF patients.^24, 25^ Additionally, monocyte-derived fibrocytes are also critical contributors to the pathogenesis of MF patients and mouse models.^26, 27^ However, specific molecular mechanisms connecting monocytes to MF are not yet fully understood.

*Jak2^V617F^* inducible-recombinant mice develop a PV phenotype with limited bone marrow fibrosis.^28^ Here we demonstrate that, *Nras^G12D/+^Jak2^V617F/+^* mice developed monocytosis and accelerated MF. In these mice, we identified that Ly6c^high^ monocytes that overexpressed *CD38* were accompanied with lower levels of nicotinamide adenine dinucleotide (NAD)^+^ and exhibited increased quantity of monocyte derived fibrocytes *in vitro* and *in vivo*. Pharmacological targeting of CD38, a NAD^+^ hydrolase, could inhibit the differentiation of fibrocytes *in vitro* and prevent fibrogenesis *in vivo*. In humans, we also observed that CD14^+^ monocytes from MF patients exhibited significantly greater levels of CD38 expression than controls. Therefore, increased expression of CD38 in monocytes may serve as a marker for the progression of fibrosis in patients with MPN, and targeting CD38 could represent a novel approach for antifibrotic therapy.

## Methods

### Patients

Peripheral blood was obtained from patients with healthy controls (HCs), Chronic myelomonocytic leukemia (CMML), PV, ET, and MF. Diagnoses were classified according to 2022 World Health Organization criteria.^29^ Studies involving human samples were approved by the Ethics Committee of the Blood Disease Hospital, Chinese Academy of Medical Sciences & Peking Union Medical College.

### CD14^+^ Monocyte Isolation

Peripheral blood mononuclear cells were isolated by density gradient, using Lymphocyte Separation Medium (TBD science). Monocytes were selected using human CD14 MicroBeads (Miltenyi Biotec) and MACS separation column following the manufacturer’s instructions.

### Mice

*Jak2^V617F/+^* mice were used as previously described.^28^ *Nras^G12D/+^* mice were kindly provided by Dr. Xiaofan Zhu (State Key Laboratory of Experimental Hematology). Their genotypes were identified by PCR at genomic DNA level (Figure S2B). For Cre induction, mice were intraperitoneally injected with 15 mg/kg polyinosinic-polycytidylic acid (poly (I: C); InvivoGen) every other day 3 times at 6-8 weeks old.

**Detailed Methods are provided in the Supplemental Materials.**

## Results

### *Nras^G12D/+^* induces an early onset of anemia and monocytosis in *Jak2^V617F/+^* mice

In a dataset including 715 MF patients (55 post-PV MF, 88 post-ET MF, 117 pre-PMF and 455 overt-PMF), there were 112 cases with monocytosis (≥ 1×10^9^/L). Patients with monocytosis had more severe fibrosis (Grade2-3: 90.2% vs 82.4%, *p*=0.042, Figure S1A) and inferior overall survival (OS) (median survival: 62 months vs 157 months, *p* < 0.0001) (Figure S1B). Patients with monocytosis were more likely to have mutated *ASXL1* (49.5% vs 25.7%; *p* < 0.001), *SETBP1* (10.8% vs 3.0%; *p* < 0.001), *SRSF2* (11.7% vs 3.5%; *p* < 0.001), *NRAS* (9.1% vs 2.5%; *p* = 0.002) and *EZH2* (12.0% vs 5.0%; *p* = 0.005) (Figure S1C). Activating mutations in *RAS*, particularly *NRAS*, are common in myeloid malignancies.^30^ MF patients harboring *RAS* or *CBL* mutations were more likely to be diagnosed with overt PMF compared to pre-PMF and were associated with inferior OS.^31^ Given the association between RAS pathway gene mutations and monocytosis and advanced fibrosis in our analysis. We elected to investigate the role of monocytosis and fibrosis by establishing a murine model with dual mutations in *Jak2* and *Nras*.

We generated *Mx1-Cre* mice, *Nras^G12D/+^*, *Jak2^V617F/+^*, and *Nras^G12D/+^Jak2^V617F/+^*mice. The first administration of the poly (I: C) injection was designated as day 1 (Figure S2A-B). We confirmed *Ras* G12D mutant protein expression in whole BM cells of *Nras^G12D/+^* and *Nras^G12D/+^Jak2^V617F/+^*mice after poly (I: C) for 2 weeks (Figure S2C). Compared with age-matched J mice, *Nras^G12D/+^Jak2^V617F/+^* mice showed higher white blood cell (WBC) and monocyte counts in peripheral blood (PB) (Figure S2D). Both *Jak2^V617F/+^* and *Nras^G12D/+^Jak2^V617F/+^* mice developed erythrocytosis and died of thrombosis at an early stage of the disease (Figure S2E-F). *Nras^G12D/+^Jak2^V617F/+^* mice showed marked splenomegaly already 6 weeks after poly (I: C) (Figure S2G). Subsequently, we conducted methylcellulose colony-forming assays and observed that BM and spleen cells from *Nras^G12D/+^Jak2^V617F/+^* mice exhibited a higher number of colonies, especially granulocyte-macrophage colony forming unit (CFU-GM) in comparison to cells from other genotypes (Figure S2H). This suggests that *Nras^G12D/+^Jak2^V617F/+^* mice possess more hematopoietic stem and progenitor cells (HSPCs) in BM, as well as a higher level of activated extramedullary hematopoiesis in spleens.

We then transplanted BM cells collected from *Mx1-Cre* mice, *Nras^G12D/+^*, *Jak2^V617F/+^*, and *Nras^G12D/+^Jak2^V617F/+^* mice into lethally irradiated wild-type mice (Figure 1A), hereafter referred to as WT, N, J, and NJ mice, respectively. Eight weeks after transplantation, PB analyses showed an increase in WBC counts, monocyte counts, and hemoglobin levels accompanied with a significant increase in the frequency of both CD11b^+^CD115^+^Ly6c^high^(classical or inflammatory monocytes^32^)and CD11b^+^CD115^+^ Ly6c^low^(non-classical or patrolling monocytes^33^) monocytes in NJ mice (Figure 1B-C, Figure S3A). Moreover, the proportion of c-kit^+^ cells in PB was significantly higher in NJ mice, with most of these cells being CD11b^+^(Figure 1D). However, the hemoglobin in NJ mice began to decline after 20 weeks of transplantation. Mice receiving NJ cells showed reduced OS compared with J (median survival: 292 days vs 340days, *p* = 0.0292) (Figure 1E). The spleen of NJ mice showed marked distortion of the normal architecture and extramedullary hematopoiesis (Figure 1F-G) at 20 and 28 weeks after transplantation. Consistent with spleen histology analysis, these mice showed an increased population of Lineage^−^Sca-1^+^c-Kit^+^ (LSK) and myeloid progenitors (Lineage^−^Sca-1^−^c-kit^+^, MP), indicating more pronounced extramedullary hematopoiesis (Figure 1H). Furthermore, flow cytometry analysis of splenocytes revealed massive infiltration by Ly6c^high^ and Ly6c^low^ monocytes in NJ mice compared with the other three groups, in keeping with PB data (Figure 1I).

**Figure 1.**
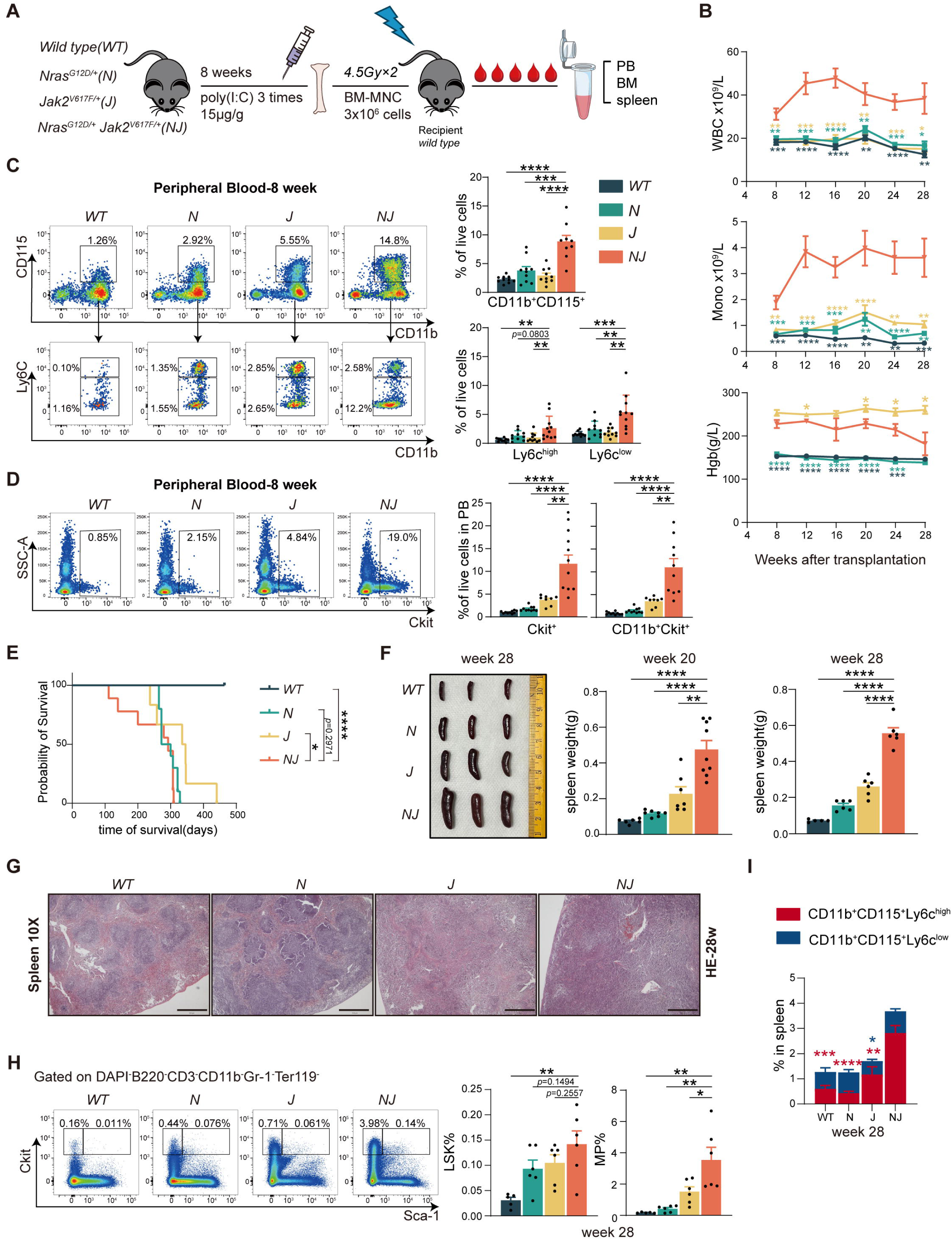
*Nras^G12D/+^* induces an early onset of anemia and monocytosis in *Jak2^V617F/+^* mice. **A.** Schematic of the transplanted *Nras^G12D/+^Jak2^V617F/+^*-Mx1-Cre mouse model. **B.** White blood cell (WBC) counts, monocyte counts and hemoglobin level of wild type (WT, *n* = 10), *Nras^G12D/+^*(N, *n* = 8), *Jak2^V617F/+^*(J, *n* = 8), and *Nras^G12D/+^Jak2^V617F/+^* (NJ, *n* = 10) recipient mice following transplantation over a 28-week period. **C-D.** Representative flow cytometry dot plot and the percentage of CD11b^+^CD115^+^ monocyte and ckit^+^ cells in bone marrow (BM). **E.** Kaplan-Meier survival curve for mice with NJ, J, N and WT. Statistics were assessed by Log-rank test. **F.** Spleen size and weight of four groups of mice at 20 weeks and 28weeks. **G.** Hematoxylin and eosin (H&E) staining of spleen of age-match (28 weeks) mice. Scale = 500 μm; Original magnification, 10×. **H.** Representative flow cytometry plots and the percentage of Lin^−^Sca1^+^c-Kit^+^ (LSK) and Lin^−^ Sca-1^−^c-kit^+^ (myeloid progenitor, MP) cells in spleen from WT, N, J, and NJ mice 28 weeks post-transplantation. **I.** The percentage of CD11b^+^CD115^+^ monocyte in spleen 28 weeks post-transplantation. Red asterisks represent the statistical comparison of Ly6c^high^ monocytes among the four groups, while blue asterisks represent the statistical comparison of Ly6c^low^ monocytes among the four groups. All data represent mean ± standard error of the mean (SEM). Statistics were assessed by two-tailed Student’s T test. \**P* ≤ .05; \*\**P* ≤ .01; \*\*\**P* ≤ .001; \*\*\*\**P* ≤ .0001.

### *Nras^G12D/+^* accelerated *Jak2^V617F/+^*-induced myelofibrosis

BM histopathology showed only light fibrosis in the NJ mice’s sternum at 20 weeks (NJ vs J: Grade 1 vs Grade 0, *p*=0.0427), but at 28 weeks, both the sternum and femur were significantly fibrotic (NJ vs J: Grade 2 vs Grade 0, *p*=0.002) (Figure 2A). In contrast, BM from J mice exhibited no fibrosis at 20 weeks and only mild fibrosis in the sternum at 28 weeks. Fibrosis was significantly accelerated in the context of monocytosis and *Nras^G12D^*co-expressed with *Jak2^V617F/+^* (Figure 2B). Considering fibrosis development in NJ mice, we divided the observation period into two phases: pre-fibrotic stage (20 weeks) and fibrotic stage (28 weeks).

**Figure 2.**
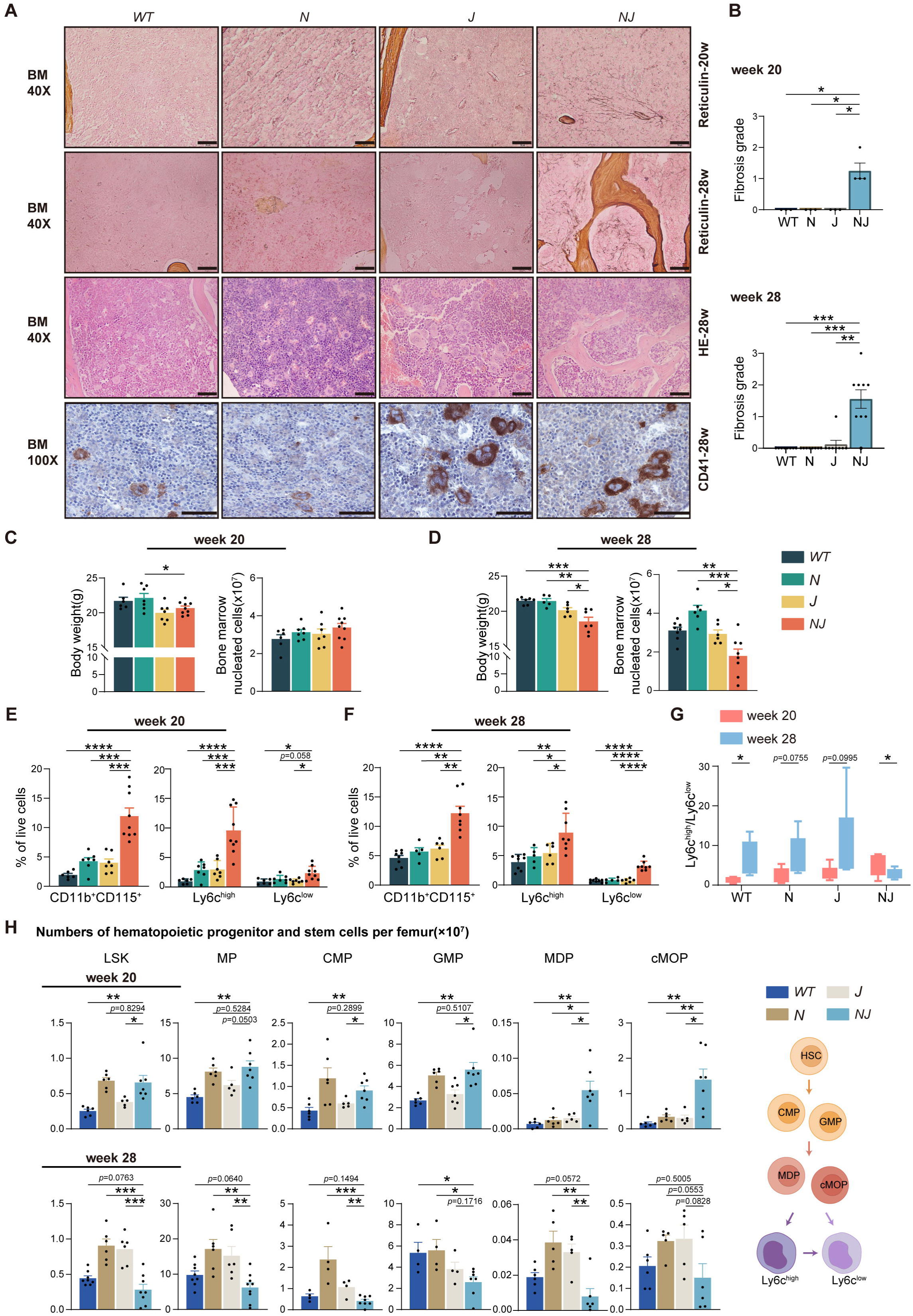
*Nras^G12D/+^* accelerates *Jak2^V617F/+^*-induced myelofibrosis. **A.** Reticulin staining of BM at 20 and 28 weeks and H&E staining at 28 weeks after transplantation. CD41-stainging of BM at 28 weeks. Scale = 50 μm; Original magnification 40× for reticulin and H&E staining and 100× for CD41-stainging. **B.** Comparison of BM fibrosis grade in NJ, J, N, and WT at 20 and 28 weeks. Statistics were assessed by Kruskal-Wallis test. **C-D.** Body weight and BM nucleated cells count per femur of four groups of mice at 20 and 28 weeks. **E-F.** The percentage of CD11b^+^CD115^+^ monocyte in BM at 20 and 28 weeks. **G.** The ratio of Ly6c^high^/Ly6c^low^ monocyte in BM at 20 and 28 weeks. **H.** The absolute number of Lin^−^Sca1^+^c-Kit^+^ (LSK) and Lin^−^ Sca1^−^ckit^+^ (myeloid progenitor, MP) cells, common myeloid progenitors (CMP, Lin^−^Sca-1^−^c-kit^+^CD34^+^FcγRII/III^low^), granulocyte-monocyte progenitors (GMP, Lin^−^Sca-1^−^c-kit^+^CD34^+^FcγRII/III^high^), monocyte-dendritic cell progenitors (MDP, Lin^−^Sca-1^−^c-kit^+^CD115^+^FcγRII/III^high^CD135^+^Ly6c^-^), and common monocyte progenitors (cMOP, Lin^−^Sca-1^−^c-kit^+^CD115^+^FcγRII/III^high^ CD135^-^ Ly6c^+^) in BM of four groups at 20 and 28 weeks. All data represent mean ± standard error of the mean (SEM). Statistics were assessed by two-tailed Student’s T test. \**P* ≤ .05; \*\**P* ≤ .01; \*\*\**P* ≤ .001; \*\*\*\**P* ≤ .0001.

Another characteristic of MPN-associated MF is the augmentation in the quantity of dysplastic megakaryocytes. NJ and J mice exhibited a substantially higher number of CD41^+^ megakaryocytes in BM than WT and N mice (Figure 2A). NJ mice also showed a modest decrease in megakaryocyte ploidy in comparison to J mice (Figure S3B). There was no notable disparity in the quantity or ploidy of megakaryocytes between N and WT mice. At the pre-fibrotic stage, NJ and J mice had lower body weights than WT and N mice, and the number of femoral cells in NJ mice had increased slightly, indicating a high proliferation phase (Figure 2C). As MPN disease advanced, NJ mice displayed decreased cellularity in femurs and weight loss (Figure 2D). At both 20 weeks and 28 weeks, NJ mice presented a significantly higher percentage of CD11b^+^CD115^+^ total monocytes and Ly6c^high^ and Ly6c^low^ subpopulation in BM (Figure 2E-F). Relative to the fibrotic stage, the increase in the CD11b^+^CD115^+^Ly6c^high^ monocytes was more obvious in the pre-fibrotic stage. The ratio of Ly6c^high^ to Ly6c^low^ was elevated in NJ mice at 20 weeks compared to the other three groups, suggesting a more pronounced inflammatory state in the BM. At 28 weeks, the ratio was significantly decreased (Figure 2G). We then analyzed HSPCs in the BM of mice. LSK and MP were substantially elevated in NJ mice during the pre-fibrotic stage, which was concurrent with an increase in downstream common myeloid progenitors (CMP), granulocyte-monocyte progenitors (GMP), monocyte-dendritic cell progenitors (MDP) and common monocyte progenitors (cMOP) (Figure 2H). The Gating strategy is shown in Figure S3C-D. These findings indicated that NJ HSPCs exhibits a differentiation bias toward the monocyte lineage. At the fibrosis stage, the number of HSPCs in NJ mice was significantly lower than that of the other three groups. Thus, the NJ mouse model demonstrates a lineage bias towards monocytosis with subsequent fibrotic progression of disease and reduction of BM cellularity and HSPC representation, consistent with disease progression.

### *Nras^G12D/+^* increased monocyte-derived fibrocyte differentiation in *Jak2^V617F/+^* mice

Several studies have identified neoplastic monocyte-derived fibrocytes as a potential contributor to MF.^26, 27^ The increased monocytes in NJ mice motivated us to investigate whether monocyte-derived fibrocytes play an important role in the formation of MF. We isolated BM cells from NJ, J, N and WT mice and cultured them in conditions that promote the differentiation of monocytes to fibrocytes.^34^ Cells derived from NJ mice’s BM nucleated cells produced a greater number of long spindle-shaped CD45^+^CollagenI^+^ fibrocytes on day 5 than cells from other three groups (Figure 3A-B). To identify from which monocyte subgroup the enhanced fibrocytes originated, we sorted CD11b^+^CD115^+^Ly6c^high^ and CD11b^+^CD115^+^Ly6c^low^ monocytes and seeded them in the fibrocyte culture medium. The Ly6c^high^ monocytes formed a greater number of fibrocytes in NJ mice whereas less fibrocytes were formed from all genotypes using Ly6c^low^ monocytes (Figure 3C-D, Figure S3E). Flow cytometry analysis of BM from NJ mice also showed markedly increased proportions of monocyte-derived fibrocytes (CD45^+^CollagenI^+^, CD11b^+^CollagenI^+^ or CD68^+^CollagenI^+^) compared with BM from other mice (Figure 3E).

**Figure 3.**
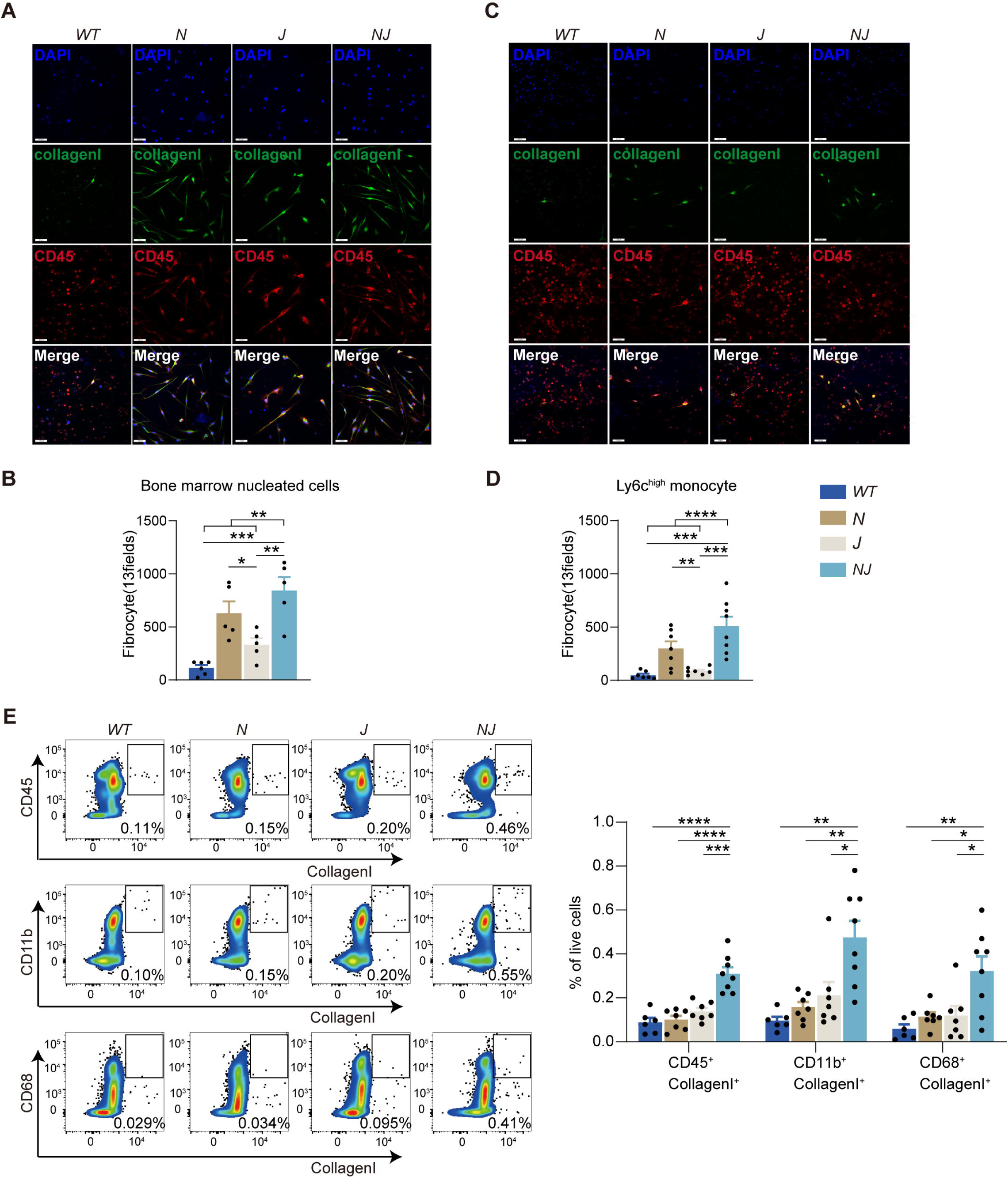
*Nras^G12D/+^* increased monocyte-derived fibrocyte differentiation in *Jak2^V617F/+^*mice. **A-B.** Representative immunofluorescence imaging and numbers of cultured monocyte-derived fibrocytes derived from BM nucleated cells among four groups. **C-D.** Representative immunofluorescence imaging and numbers of cultured monocyte-derived fibrocytes derived from Ly6c^high^ monocyte among four groups. Scale = 40 μm; Original magnification 20×. **E.** Representative flow cytometry plots and the percentage of monocyte-derived fibrocytes among four groups. All data represent mean ± standard error of the mean (SEM). Statistics were assessed by two-tailed Student’s T test. \**P* ≤ .05; \*\**P* ≤ .01; \*\*\**P* ≤ .001; \*\*\*\**P* ≤ .0001.

Altogether, these data indicate that the increased neoplastic monocyte-derived fibrocytes may be linked to the hastening of BM fibrosis in NJ mice. Fibrocyte mainly differentiate from the Ly6c^high^ monocyte subpopulation. In N mice, despite exhibiting a potential capacity for fibrocyte generation *in vitro*, MF did not occur *in vivo* with relatively fewer fibrocytes detected, indicating cooperative with *Jak2^V617F^* is necessary to produce this phenotype.

### Mult-omics profiling revealed significantly elevated expression of CD38 and abnormal NAD^+^ metabolism in the *Nras^G12D/+^ Jak2^V617F/+^* Ly6c^high^ monocytes

To elucidate the molecular mechanism of MF, we performed bulk RNA sequencing on NJ, J, N and WT mice on sorted CD11b^+^CD115^+^Ly6c^high^ cells (the primary origin of monocyte-derived fibrocytes) at pre-fibrotic stage to elucidate the transcriptional alterations between genotypes.

Comparison of differentially expressed genes (fold change ≥ 1-fold; *p* ≤ 0.05) in Ly6c^high^ monocyte revealed 229 genes with altered expression between NJ mice and other genotypes (Figure 4A, Figure S4A) (99 were downregulated and 130 were upregulated). Analysis of Kyoto Encyclopedia of Genes and Genomes (KEGG) pathways revealed the enrichment of hematopoietic cell lineage and several inflammation-related pathways including NF-κB and TNF pathways (Figure 4B). Given that N mice exhibit enhanced fibrocyte formation *in vitro* but do not manifest fibrosis *in vivo*, our investigation primarily focused on the differences between NJ and N mice. Compared with N mice, monocytes of NJ mice showed significantly elevated expression of *Cd38* (Figure 4C). The results were consistent with those observed in WT and J mice (Figure S4B). We validated *Cd38* upregulation in NJ Ly6c^high^ monocytes and fibrocytes at the mRNA level using quantitative reverse transcriptase polymerase chain reaction(qRT-PCR) (Figure 4D) in independent cohorts. We hypothesized that monocytes that overexpress CD38 have strong potential to differentiate to fibrocytes and may accelerate MF. Our group previously found that *Asxl1* mutations accelerate MF in the context of *Jak2^V617F^* via EGR1-TNFA axis.^34^ Here, we also found that significant increased *Cd38* mRNA level in Ly6c^high^ monocytes of *Asxl1^-/-^ Jak2^V617F^* mice than *Jak2^V617F^*mice (Figure S4C). This discovery supports our hypothesis regarding the potential role of CD38 in the progression of MF.

**Figure 4.**
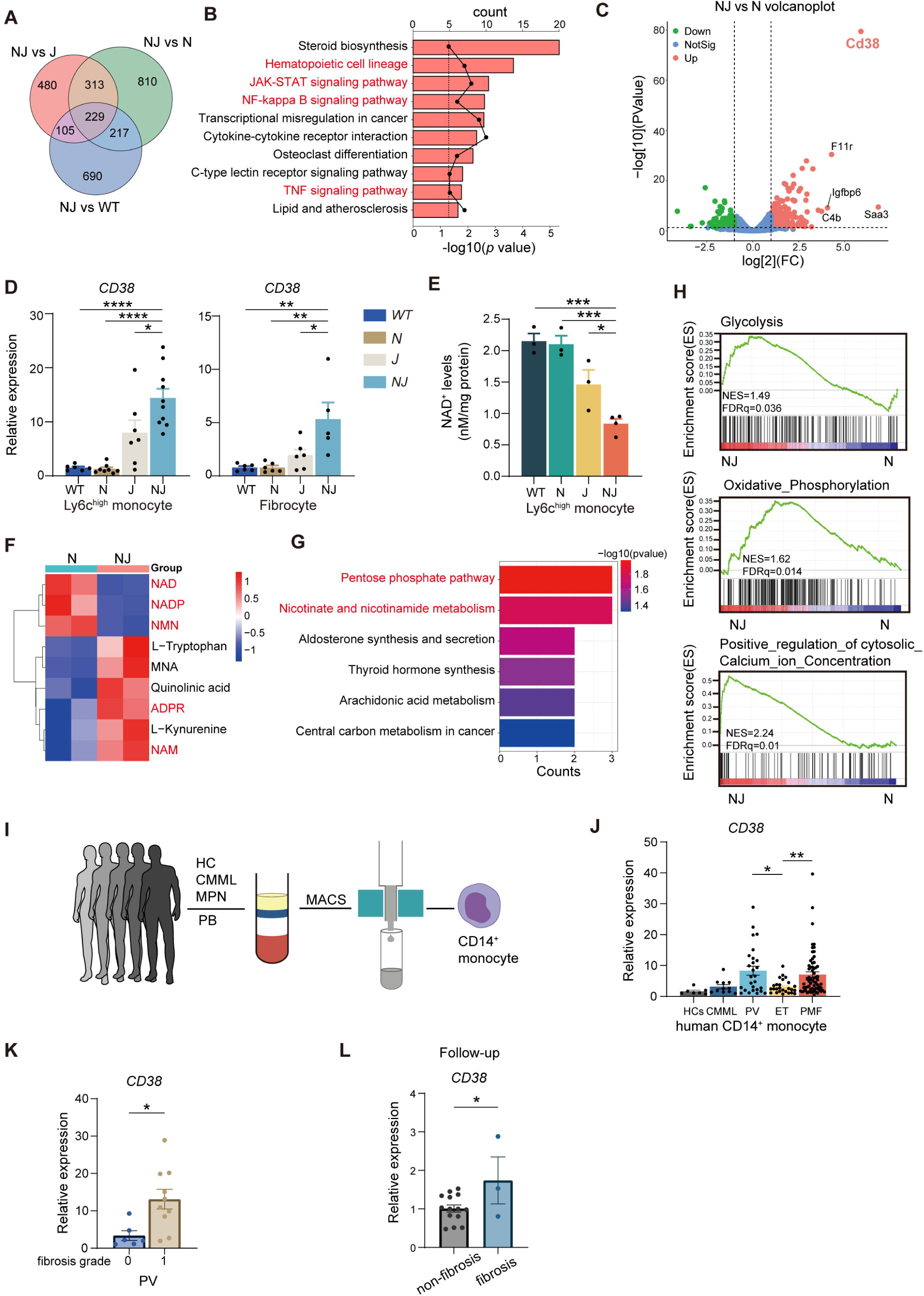
Mult-omics sequencing revealed significantly elevated expression of CD38 and abnormal NAD^+^ metabolism in the *Nras^G12D/+^ Jak2^V617F/+^* Ly6c^high^ monocytes. A. Venn diagram of differentially expressed genes (DEGs) in NJ compared with J, N and WT BM CD11b^+^CD115^+^Ly6c^high^ monocyte (fold change≥1 fold; *p*≤ .05). **B.** Kyoto Encyclopedia of Genes and Genomes (KEGG) pathway enrichment analysis of DEGs. **C.** Volcano plot illustrates DEGs between NJ and N. **D.** Quantification of *Cd38* messenger RNA (mRNA) levels in sorted BM CD11b^+^CD115^+^Ly6c^high^ monocyte and cultured monocyte-derived fibrocytes from WT, N, J and NJ 20 weeks post-transplantation by qRT-PCR. **E.** The NAD^+^ levels in CD11b^+^CD115^+^Ly6c^high^ monocyte from WT, N, J and NJ at 20 weeks post-transplantation. **F.** Heatmap illustrates the relative abundance of metabolites in the NAD^+^ metabolic pathway in NJ compared to N. **G.** KEGG pathway enrichment analysis based on the altered metabolites identified in NJ compared to N. **H.** Gene set enrichment analysis (GSEA) showed that enrichment gene sets between NJ and N. **I.** Schematic representation of experimental procedures. **J.** Quantification of mRNA levels of *CD38* in PB CD14^+^ monocyte; HCs (*n* = 7), CMML (*n* = 11), PV (*n* = 27); ET (*n* = 26); PMF (*n* = 65). **K.** Comparison of *CD38* expression between PV with BM fibrosis grade 0(*n* = 6) and grade 1(*n* = 11). **L.** Comparison of baseline CD38 expression between MPN patients who developed fibrosis (n = 3) and those who did not (n = 14). NAD^+^: nicotinamide adenine dinucleotide; NADP: Nicotinamide adenine dinucleotide phosphate; NMN: Nicotinamide mononucleotide; MNA: methylnicotinamide; ADPR: ADP-ribose; NAM: nicotinamide. All data represent mean ± standard error of the mean (SEM). Statistics were assessed by two-tailed Student’s T test. \**P* ≤ .05; \*\**P* ≤ .01; \*\*\**P* ≤ .001; \*\*\*\**P* ≤ .0001.

CD38, a type II transmembrane nicotinamide adenine dinucleotide (NAD)^+^ hydrolase, governs cellular processes by converting NAD^+^ into nicotinamide (NAM) and ADP-ribose (ADPR) or cyclic ADP-ribose (cADPR), which act as second messengers to mediate calcium signaling.^35–37^ As a key regulator of NAD^+^ bioavailability, CD38 critically impacts energy metabolism pathways including glycolysis and oxidative phosphorylation through its enzymatic activity.^38^ Considering the significant overexpression of *Cd38* in NJ Ly6c^high^ monocytes, we tested the NAD^+^ level in mice with different genotypes. We found that NJ Ly6c^high^ monocytes had significantly lower NAD^+^ levels (Figure 4E). As NAD^+^ levels are determined by a balance between synthesis and degradation, we measured the mRNA levels of the enzymes that were involved in the biosynthesis or metabolism of NAD^+^. The mRNA levels of enzymes in the de novo NAD-synthetic pathway and NAM-salvage-pathway enzymes exhibited no variation among the four groups. Among NAD hydrolases, only *Cd38* expression increased significantly in NJ mice (an 9.6-fold increase compared to WT), while the others showed no significant changes (Figure S4E).

Quasi-Targeted Metabolomics demonstrated an abnormal NAD^+^ metabolism with a significant reduction in NAD^+^ levels and nicotinamide mononucleotide (NMN, the NAD^+^ precursor), accompanied by an elevation in its metabolic derivatives, NAM and ADPR in NJ Ly6c^high^ monocytes compared to N mice, as shown in Figure 4F. NAD^+^ is essential for both glycolysis and oxidative phosphorylation, accepting electrons from glycolysis and feeding them to oxidative phosphorylation that declines during aging.^39^ From both mRNA and metabolite profiling, pathway enrichment analysis revealed that pathways related to glycolysis, oxidative phosphorylation, and calcium concentration were positively enriched in NJ monocytes compared with N(Figure 4G-H).

We then examined the expression of CD38 in patients with MPNs. As our N mice develop chronic myelomonocytic leukemia (CMML)-like phenotype, patients with CMML were also included in the study (Figure 4I). By qRT-PCR analysis, we found CD14^+^ monocytes (equivalent to mouse classical monocytes) from PV and PMF showed significantly higher mRNA expression of *CD38*, compared with healthy controls (HCs), CMML, or ET (Figure 4J). Monocytes from PV patients with grade 1 fibrosis had higher *CD38* expression than those without fibrosis (Figure 4K). Furthermore, a retrospective analysis of the CD38 expression in BM CD14^+^ monocytes from PV or ET patients at first diagnosis. Our data revealed that BM CD14^+^ monocytes from patients who progressed to post-PV or ET MF during following up exhibited significantly higher baseline CD38 expression compared to those who did not develop fibrosis (Figure 4L). We also investigated alterations in other NAD^+^ synthases and NAD^+^-consuming enzymes in CD14^+^ monocytes. Interestingly, the expression of the enzyme IDO1 was significantly increased in monocytes of PV patients, suggesting that these cells may be protected from the high *CD38* expression by increasing NAD^+^ levels via the de novo pathway (Figure S4F). In summary, these data suggest that elevated CD38 expressions in monocytes may act as a biomarker for the progression to MF.

### Overexpression of CD38 in monocytes induced bone marrow fibrosis

It has been reported that CD38 expression is regulated at the transcriptional level by stimulation from TNF and the NF-κB pathways.^40^ TRRUST (Transcriptional Regulatory Relationships Unraveled by Sentence-based Text mining) database revealed that *Cd38* may be regulated by nuclear factor κ light-chain enhancer of activated B cells (Nfkb1) transcription factor (Figure S5A). To investigate whether inflammatory cytokines in BM microenvironment responsible for the rise in *Cd38* expression in monocytes, we administered several cytokines to CD115^+^ monocytes. TNF-α and IL-6 can substantially stimulate the increase in monocyte CD38 expression (Figure S5B), while IL-1β requires higher concentrations to achieve a similar effect. This is consistent with previous results^41^. Conversely, greater concentrations of Tgfβ were unable to stimulate its increase. Serum analysis across the four genotypes mice revealed a significant increase in TNF-α levels at both 20 and 28 weeks (Figure S5C). We also found a positive correlation between the expression of *CD38* and serum TNF-α levels in MPN patients (Figure S5D). We then used TNF-α to stimulate CD14^+^ monocytes from PV or ET patients and found it can also induce the upregulation of *CD38*(Figure S5E). These results align with the findings from RNA sequencing, which showed upregulation of the TNF-NF-κB signaling pathway in NJ Ly6c^high^ monocytes (Figure S4D).

As lipopolysaccharide (LPS) can greatly increase Cd38 mRNA levels in macrophages,^41, 42^ we isolated BM CD115^+^ monocytes from N mice and treated them with LPS for 100 ng/ml for 16 hours. The expression of CD38 on monocytes can be increased by 80.42-fold in response to LPS (relative to PBS) (Figure S5F). Considering that N mice produce more fibrocytes *in vitro* but do not develop fibrosis *in vivo*, we tested whether increasing monocyte *Cd38* expression by intraperitoneal injection of LPS could induce fibrosis in N mice. We treated N mice for 4 weeks with a chronic sublethal dose of LPS (Figure S6A). Three days after the last dosing, mice received LPS showed significant anemia compared with those receiving PBS (Figure S6B). BM histopathology showed significant deposition of reticular fibers and BM sclerosis in the femur of LPS group (Figure S6C). In contrast, no fibrosis was observed in the PBS group. Furthermore, N mice in the LPS group exhibited a significant decrease in the quantity of femoral BM cells, together with the presence of little extramedullary hematopoiesis (slightly enlarged spleen) (Figure S6D-E). QRT-PCR analysis revealed a substantial rise in the mRNA levels of *Cd38* in monocytes in BM Ly6c^high^ monocytes of LPS group (Figure S6F). Flow cytometry analysis revealed a notable increase in monocyte-derived fibrocytes in BM of LPS group (Figure S6G-H).

This indicates that N mice exhibit a greater intrinsic tendency for fibrosis and further underscores the pivotal role of *CD38* overexpressed monocytes in the pathophysiology of MF.

### The CD38 inhibitor 78c delayed the onset of myelofibrosis by restoring monocyte NAD^+^ homeostasis

In view of the substantial fibrosis progression accompanied by elevated CD38 expression and a decline of NAD^+^ levels that was noted in multiple organ fibrosis,^43, 44^ Metabolomics also demonstrated a significant reduction in NAD^+^ levels and NMN, the NAD^+^ precursor in NJ mice. These indicate a correlation between disruption in NAD^+^ metabolism in monocyte and fibrogenesis. Therefore, we investigate whether boosting NAD^+^ levels by selectively inhibiting its CD38-mediated hydrolysis by 78c (a highly potent and specific thiazoloquin(az)olin(on)e-based CD38 Inhibitor with cell permeability and dual inhibitory activity on ecto- and intracellular CD38 functions^45, 46^) or supplementing NAD^+^ precursors NMN can alleviate MF. We incubated NJ BM nucleated cells with increasing concentrations of 78c and NMN and assessed disparities in the quantity of fibrocytes. We discovered that both 78c and NMN could reduce the generation of fibrocytes and increase the NAD^+^ levels in CD115^+^ monocytes in a dose-dependent manner (Figure 5A-B). Given the absence of any variation in the apoptosis rate of BM cells following a 24-hour exposure to 78c and NMN (Figure S7A-B), we hypothesized that replenishing NAD^+^ levels can impede the differentiation of monocytes into fibrocytes. We incubated PB CD14^+^ monocytes of MF patients in culture under conditions that induce differentiation to fibrocytes with 78c and NMN. Both 78c and NMN restored NAD^+^ level and inhibited the differentiation of fibrocytes (Figure 5C-Dd). These results suggest that restoring NAD^+^ levels can inhibit the differentiation of monocytes into fibrocytes.

**Figure 5.**
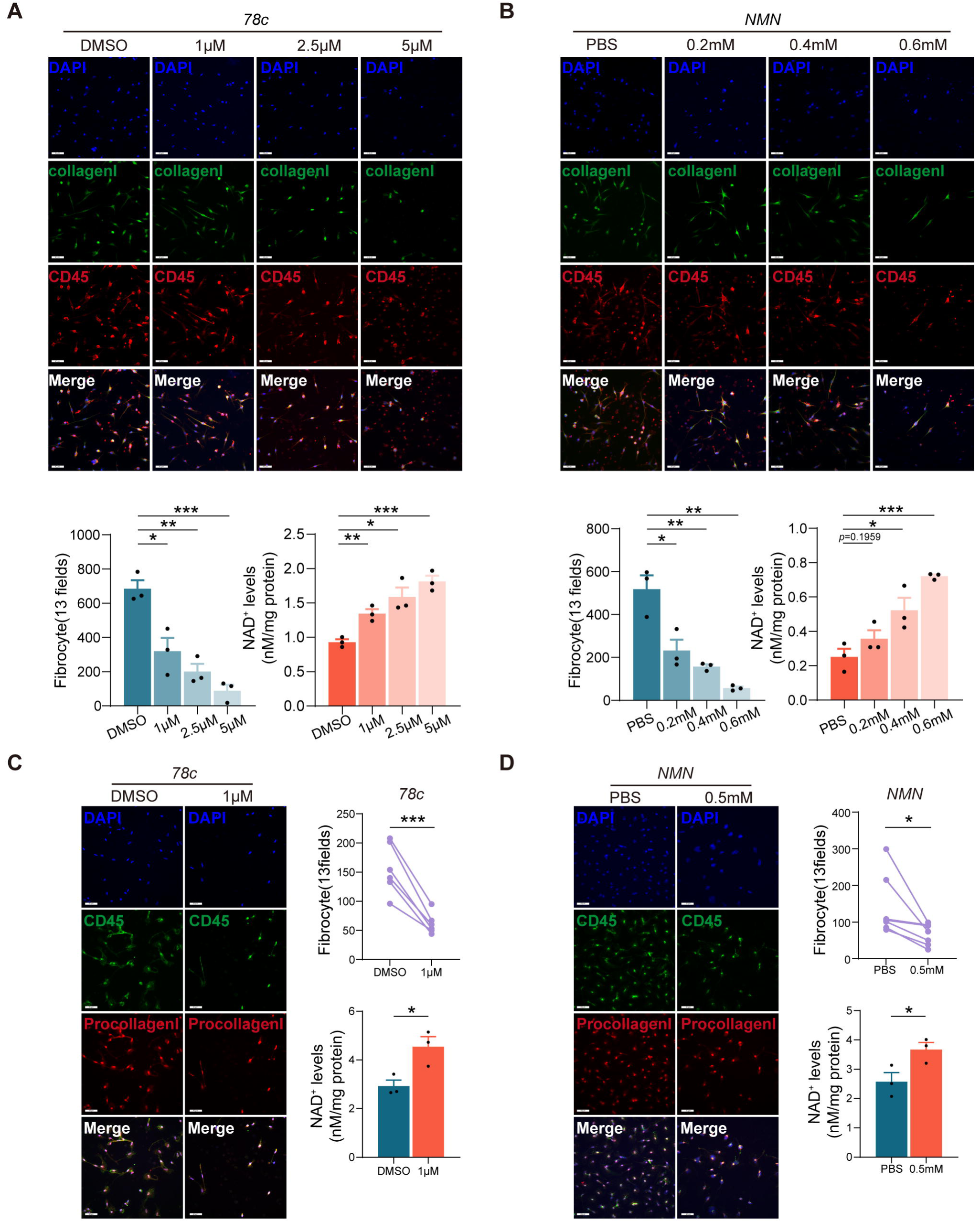
78c and NMN inhibited differentiation of monocytes to fibrocyte by restoring monocyte NAD^+^ homeostasis. **A-B.** Representative immunofluorescence imaging (upper) and numbers (lower) of cultured fibrocytes derived from NJ BM nucleated cells treated with different concentrations of 78c and NMN. The NAD^+^ levels in CD115^+^ monocytes were evaluated after exposure to 78c and NMN for 16 hours *in vitro*. Scale = 40 μm; Original magnification 20×. **C-D.** Representative immunofluorescence imaging (left) and counts (right) of monocyte-derived fibrocytes derived from MF patients’ PB CD14^+^monocyte treated with 78c and NMN. The NAD^+^ levels in CD14^+^ monocytes were evaluated after exposure to 78c and NMN for 16 hours *in vitro*. Scale = 40 μm; Original magnification 20×. All data represent mean ± standard error of the mean (SEM). Statistics were assessed by two-tailed Student’s T test or Mann-Whitney U test. \**P* ≤ .05; \*\**P* ≤ .01; \*\*\**P* ≤ .001; \*\*\*\**P* ≤ .0001.

To examine the effect of 78c *in vivo*, NJ mice were randomly allocated into two groups that received 4-week solvent (Control, n=6) or 78c (n=5) treatment 20 weeks after transplantation (Figure 6A). After a 4-week follow-up, control group mice exhibited a reduction in the levels of WBC, monocytes, and hemoglobin. Conversely, the 78c group mice maintained a high Hgb level (Figure 6B) and prevented the progressive anemia typically seen in NJ mice. BM histology revealed a marked reduction in reticular fibers (Figure 6C-D). *Cd38* mRNA level in BM Ly6c^high^ monocytes was significantly downregulated in 78c group (Figure 6E).

**Figure 6.**
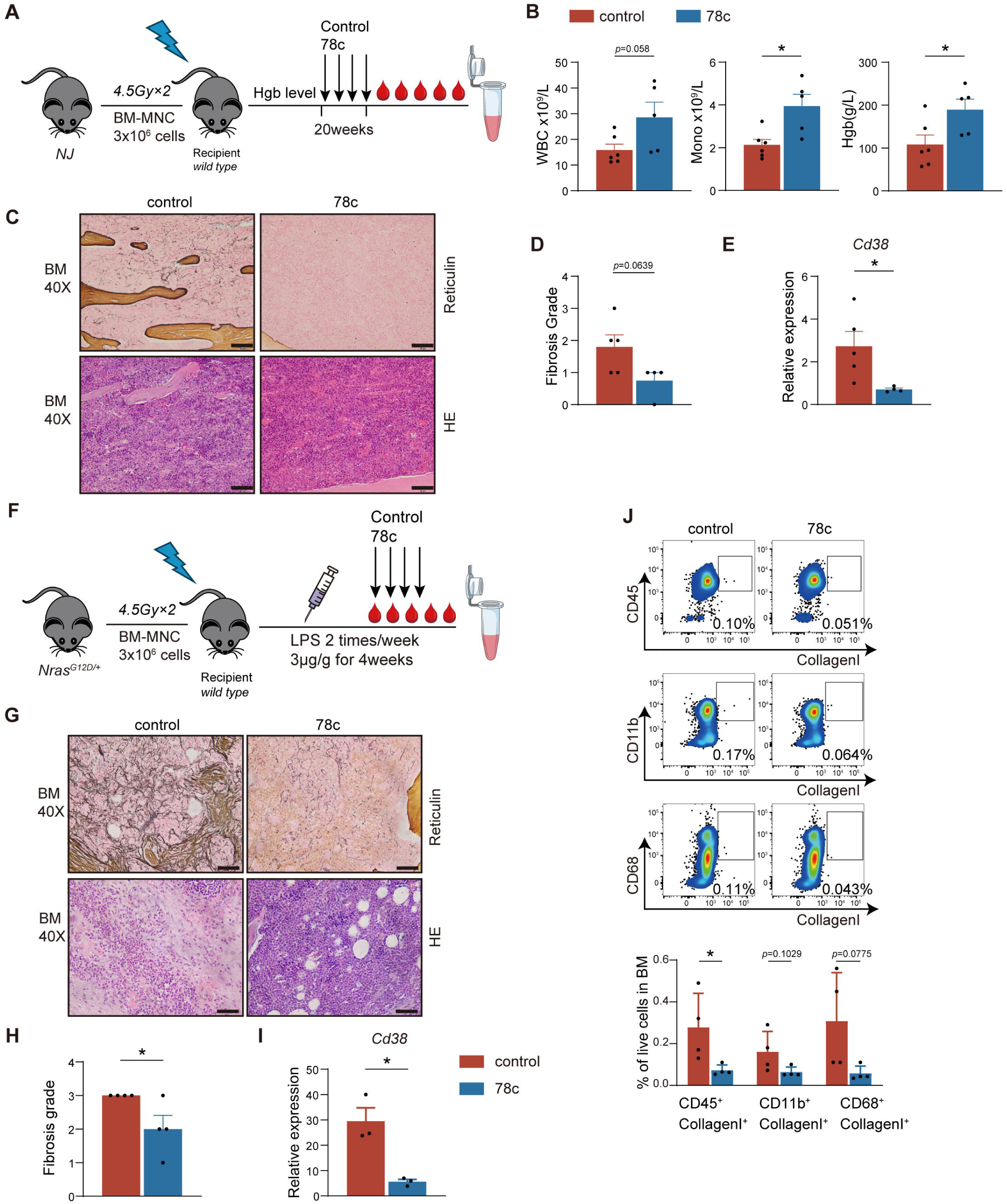
CD38 inhibitor 78c delayed the onset of myelofibrosis by restoring monocyte NAD^+^ homeostasis. **A.** Schematic of the *Nras^G12D/+^Jak2^V617F/+^* mouse model treated with 78c. **B.** White blood cell (WBC) counts, monocyte counts and hemoglobin level of two groups. **C.** H&E and Reticulin staining of bone marrow (BM). Scale = 50 μm; Original magnification 40×. **D.** Comparison of BM fibrosis grade in each group. **E.** Quantification of mRNA levels of *Cd38* in sorted BM CD11b^+^ CD115^+^ Ly6c^high^ monocyte by qRT-PCR. **F.** Schematic of LPS-induced *Nras^G12D/+^*-Mx1-Cre transplanted mouse model induced treated with either solvent or 78c. **G.** H&E and Reticulin staining of BM. Scale = 50 μm; Original magnification 40×. **H.** Comparison of BM fibrosis grade in each group. **I.** Quantification of mRNA levels of Cd38 in sorted BM CD11b^+^ CD115^+^ Ly6c^high^ monocyte by qRT-PCR. **J.** The representative flow cytometry dot plot and the percentage of fibrocytes of two groups. All data represent mean ± standard error of the mean (SEM). Statistics were assessed by two-tailed Student’s T test. \**P* ≤ .05; \*\**P* ≤ .01; \*\*\**P* ≤ .001; \*\*\*\**P* ≤ .0001.

Given that LPS induces fibrosis by upregulating *Cd38* expression of monocytes in N mice, we investigated whether 78c could prevent MF onset in this model (Figure 6F). After four weeks of treatment, mice in the 78c group showed higher hemoglobin levels compared to the control group (Figure S7C). BM histology staining revealed a marked reduction in reticular fibers (Figure 6G-H). QRT-PCR further confirmed a substantial decrease in *Cd38* mRNA levels in BM Ly6c^high^ monocytes of 78c group (Figure 6I). Additionally, the proportion of monocyte-derived fibrocytes was significantly reduced (Figure 6J). These findings indicate that the inhibition of CD38 by 78c effectively attenuates the progression of MF.

## Discussion

In the pathogenesis of MPN, monocytes serve as the principal source of increased cytokines in MF patients and significantly contribute to the development of MF via monocyte-derived fibrocytes.^24, 26, 27, 34^ In MPN patients, monocytosis associates with a reduced myelofibrosis-free survival (MFS) in PV and short survival in PMF.^47, 48^ Here, consistent with those clinical observation, we have developed a genomically and biologically accurate model of fibrosis progression of MPN characterized by significantly increased counts of Ly6c^high^ inflammatory monocytes that overexpress CD38 and lose NAD^+^ metabolic hemostasis.

TNF-α and its downstream signaling pathway greatly contribute to the persistent inflammation in MPN.^49, 50^ In *Asxl1*^-/-^/*Jak2^V617F^* mice, we identified that TNF-α promoted the differentiation of monocyte-derived fibrocyte.^34^Previous data have showed that TNF-α and its downstream NF-κB signaling pathway upregulated the expression of CD38, because the *CD38* enhancer region contains two NF-κB binding sites.^51,44^ In the context of two MPN-associated MF mouse models and patients *in vivo* and *in vitro*, we identified the positive correlation between TNF-α and CD38 expression in monocytes. This data provide clear evidence that TNF-α promote the differentiation of monocyte derived fibrocyte via upregulating CD38 expression in monocytes. It provides further evidence for the fibrogenesis effect of TNF-α, which will need to be investigated in other myeloid neoplasms, such as myelodysplastic neoplasms (MDS) with BM fibrosis, using similar approaches. The differentiation of monocyte-derived fibrocyte bias play a critical role in the fibrotic formation and progression.^26, 27^ Our data suggests the differentiation form monocyte to fibrocyte induced by significantly reduced level of NAD^+^ in inflammatory monocytes due to overexpression of CD38. CD38 catalyzes the conversion of NAD^+^ into nicotinamide (NAM) and ADP-ribose (ADPR). NAD^+^, a crucial cofactor involved in regulating energy metabolism including glycolysis, and oxidative phosphorylation^38^, can also be converted into cyclic ADP-ribose (cADPR) by CD38. Both ADPR and cADPR function as second messengers, mediating various cellular processes through the mobilization of calcium ions (Ca^2+^).^36, 37^ Overexpression of CD38 participates in several pathological conditions of organic fibrosis via its enzymatic function to decrease intracellular NAD^+^ levels, such as in idiopathic pulmonary fibrosis (IPF)^52^, systemic sclerosis^43^ and renal fibrosis^44^. Here, we reported the abnormal CD38-NAD^+^ metabolic axis also took part in the pathogenesis of MF in MPN according to the multi-omics sequencing. More importantly, we identified that pharmacological inhibition by CD38 inhibitor 78c or supplementation of NAD^+^ precursor NMN enhanced NAD^+^-dependent processes and mitigate fibrosis in MPN. CD38 can locate both in the cellular and intracellular membranes and facilitate the circulation of NAD^+^ and NAD^+^ precursors both inside and outside of the cell. Whether the flux of NAD^+^ and NAD^+^ precursors in BM microenvironment promote the proliferation of mesenchymal stromal cells (MSC)-derived fibroblast (another major contributor for bone marrow fibrosis^53, 54^) remains to be investigated.

In summary, we confirmed the significant contribution of CD38-overexpressing monocytes in MF by using different MPN-associated MF murine models and retrospective analysis in MPN patients. Restoring intracellular NAD^+^ levels using the inhibitor 78c could gear down the development of MF. CD38 inhibition may serve as a therapeutic strategy to relieve bone marrow fibrosis in the clinical context. Moreover, a prospective study is urgently warranted to evaluate whether the expression of CD38 in monocytes could be a novel biomarker to precisely predict the progression of MF for PV or ET patients, even for other myeloid neoplasms.

## Acknowledgments

The authors acknowledge Xiaofan Zhu (State Key Laboratory of Experimental Hematology) for providing *Nras^G12D/+^* mice. We thank the State Key Laboratory of Experimental Hematology Flow Cytometry Center, Experimental Animal Center and Image Center for assistance with the experiments.

## Authorship Contributions

B.L., Z.X., R.K.R. and G.H. conceived the idea of this study; Y.Y. and J.L. designed the research; Y.Y., J.L., S.Z., F.L. and L.Y. performed experiments. S.Z., F.L., Z.X. and T.Q. acquired the clinical data; Y.Y. and J.L. performed statistical and bioinformatic analyses; Y.Y., X. Z., W.A., G.H., R.K.R., Z.X. and B.L. wrote the manuscript. All authors reviewed and approved the manuscript.

## Disclosure of Conflicts of Interest

R.K.R. has received consulting fees from Constellation, Incyte, Celgene/BMS, Novartis, Promedior, CTI, Jazz Pharmaceuticals, Blueprint, Stemline, Galecto, PharmaEssentia, AbbVie, Sierra Oncology, and Disc Medicines; and research funding from Incyte, Constellation, and Stemline. The remaining authors declare no competing financial interests.

## Funding information

Supported in part by CAMS Innovation Fund for Medical Sciences (Nos. 2023-I2M-2-007 and 2022-I2M-1-022), National Natural Science Foundation of China (Nos. 82170139, 82070134 and 81530008), Clinical research fund from National Clinical Research Centre for Blood Diseases (Nos. 2023NCRCA0117 and 2023NCRCA0103) and Tianjin Municipal Science and Technology Commission Grant (Nos. 23JCYBJC00480).

## Data availability statement

The data that support the findings of this study from the corresponding author upon reasonable request.

## Patient consent statement

The patients gave written informed consent compliant with the Declaration of Helsinki.

## References

1. Urrutia S, Kantarjian HM, Ravandi-Kashani F, et al. Outcomes of patients with acute myeloid leukemia and bone marrow fibrosis. J Hematol Oncol. 2024;17(1):112.

2. Zhang H, Guo W, Wang J, et al. Impact of bone marrow fibrosis on outcomes of allogeneic hematopoietic stem cell transplantation in acute myeloid leukemia. Bone Marrow Transplant. 2024;59(12):1654–66.

3. Melody M, Al Ali N, Zhang L, et al. Decoding Bone Marrow Fibrosis in Myelodysplastic Syndromes. Clin Lymphoma Myeloma Leuk. 2020;20(5):324–8.

4. Jain AG, Zhang L, Bennett JM, Komrokji R. Myelodysplastic Syndromes with Bone Marrow Fibrosis: An Update. Ann Lab Med. 2022;42(3):299–305.

5. Tefferi A. Primary myelofibrosis: 2023 update on diagnosis, risk-stratification, and management. Am J Hematol. 2023;98(5):801–21.

6. Barosi G, Mesa RA, Thiele J, et al. Proposed criteria for the diagnosis of post-polycythemia vera and post-essential thrombocythemia myelofibrosis: a consensus statement from the International Working Group for Myelofibrosis Research and Treatment. Leukemia. 2008;22(2):437–8.

7. Tefferi A, Saeed L, Hanson CA, Ketterling RP, Pardanani A, Gangat N. Application of current prognostic models for primary myelofibrosis in the setting of post-polycythemia vera or post-essential thrombocythemia myelofibrosis. Leukemia. 2017;31(12):2851–2.

8. Thiele J, Kvasnicka HM, Facchetti F, Franco V, van der Walt J, Orazi A. European consensus on grading bone marrow fibrosis and assessment of cellularity. Haematologica. 2005;90(8):1128–32.

9. Guglielmelli P, Lasho TL, Rotunno G, et al. MIPSS70: Mutation-Enhanced International Prognostic Score System for Transplantation-Age Patients With Primary Myelofibrosis. J Clin Oncol. 2018;36(4):310–8.

10. Tefferi A, Guglielmelli P, Lasho TL, et al. MIPSS70+ Version 2.0: Mutation and Karyotype-Enhanced International Prognostic Scoring System for Primary Myelofibrosis. J Clin Oncol. 2018;36(17):1769–70.

11. Verstovsek S, Mesa RA, Gotlib J, et al. A double-blind, placebo-controlled trial of ruxolitinib for myelofibrosis. New Engl J Med. 2012;366(9):799–807.

12. Harrison C, Kiladjian JJ, Al-Ali HK, et al. JAK inhibition with ruxolitinib versus best available therapy for myelofibrosis. New Engl J Med. 2012;366(9):787–98.

13. Oh ST, Verstovsek S, Gupta V, et al. Changes in bone marrow fibrosis during momelotinib or ruxolitinib therapy do not correlate with efficacy outcomes in patients with myelofibrosis. EJHaem. 2024;5(1):105–16.

14. Tefferi A, Vaidya R, Caramazza D, Finke C, Lasho T, Pardanani A. Circulating interleukin (IL)-8, IL-2R, IL-12, and IL-15 levels are independently prognostic in primary myelofibrosis: a comprehensive cytokine profiling study. J Clin Oncol. 2011;29(10):1356–63.

15. Vaidya R, Gangat N, Jimma T, et al. Plasma cytokines in polycythemia vera: phenotypic correlates, prognostic relevance, and comparison with myelofibrosis. Am J Hematol. 2012;87(11):1003–5.

16. Dunbar AJ, Kim D, Lu M, et al. CXCL8/CXCR2 signaling mediates bone marrow fibrosis and is a therapeutic target in myelofibrosis. Blood. 2023;141(20):2508–19.

17. Rai S, Grockowiak E, Hansen N, et al. Inhibition of interleukin-1β reduces myelofibrosis and osteosclerosis in mice with JAK2-V617F driven myeloproliferative neoplasm. Nat Commun. 2022;13(1):5346.

18. Rahman MF, Yang Y, Le BT, et al. Interleukin-1 contributes to clonal expansion and progression of bone marrow fibrosis in JAK2V617F-induced myeloproliferative neoplasm. Nat Commun. 2022;13(1):5347.

19. Di Buduo CA, Abbonante V, Marty C, et al. Defective interaction of mutant calreticulin and SOCE in megakaryocytes from patients with myeloproliferative neoplasms. Blood. 2020;135(2):133–44.

20. Zingariello M, Martelli F, Ciaffoni F, et al. Characterization of the TGF-β1 signaling abnormalities in the Gata1low mouse model of myelofibrosis. Blood. 2013;121(17):3345–63.

21. Woods B, Chen W, Chiu S, et al. Activation of JAK/STAT Signaling in Megakaryocytes Sustains Myeloproliferation In Vivo. Clin Cancer Res. 2019;25(19):5901–12.

22. He F, Laranjeira AB, Kong T, et al. Multiomic profiling reveals metabolic alterations mediating aberrant platelet activity and inflammation in myeloproliferative neoplasms. J Clin Invest. 2024;134(3).

23. Gangat N, Marinaccio C, Swords R, et al. Aurora Kinase A Inhibition Provides Clinical Benefit, Normalizes Megakaryocytes, and Reduces Bone Marrow Fibrosis in Patients with Myelofibrosis: A Phase I Trial. Clin Cancer Res. 2019;25(16):4898–906.

24. Fisher DAC, Miner CA, Engle EK, et al. Cytokine production in myelofibrosis exhibits differential responsiveness to JAK-STAT, MAP kinase, and NFκB signaling. Leukemia. 2019;33(8):1978–95.

25. He F, Lin S, Kong T, et al. Monocyte-Driven Aberrant Inflammation in Myeloproliferative Neoplasms Is Regulated By Galectin-1. Blood. 2024;144:875.

26. Ozono Y, Shide K, Kameda T, et al. Neoplastic fibrocytes play an essential role in bone marrow fibrosis in Jak2V617F-induced primary myelofibrosis mice. Leukemia. 2021;35(2):454–67.

27. Verstovsek S, Manshouri T, Pilling D, et al. Role of neoplastic monocyte-derived fibrocytes in primary myelofibrosis. J Exp Med. 2016;213(9):1723–40.

28. Mullally A, Lane SW, Ball B, et al. Physiological Jak2V617F expression causes a lethal myeloproliferative neoplasm with differential effects on hematopoietic stem and progenitor cells. Cancer Cell. 2010;17(6):584–96.

29. Khoury JD, Solary E, Abla O, et al. The 5th edition of the World Health Organization Classification of Haematolymphoid Tumours: Myeloid and Histiocytic/Dendritic Neoplasms. Leukemia. 2022;36(7):1703–19.

30. Bos JL. ras oncogenes in human cancer: a review. Cancer Res. 1989;49(17):4682–9.

31. Coltro G, Rotunno G, Mannelli L, et al. RAS/CBL mutations predict resistance to JAK inhibitors in myelofibrosis and are associated with poor prognostic features. Blood Adv. 2020;4(15):3677–87.

32. Sunderkötter C, Nikolic T, Dillon MJ, et al. Subpopulations of mouse blood monocytes differ in maturation stage and inflammatory response. J Immunol. 2004;172(7):4410–7.

33. Auffray C, Fogg D, Garfa M, et al. Monitoring of blood vessels and tissues by a population of monocytes with patrolling behavior. Science. 2007;317(5838):666–70.

34. Shi Z, Liu J, Zhao Y, et al. ASXL1 mutations accelerate bone marrow fibrosis via EGR1-TNFA axis mediated neoplastic fibrocyte generation in myeloproliferative neoplasms. Haematologica. 2022.

35. Piedra-Quintero ZL, Wilson Z, Nava P, Guerau-de-Arellano M. CD38: An Immunomodulatory Molecule in Inflammation and Autoimmunity. Front Immunol. 2020;11.

36. Hogan KA, Chini CCS, Chini EN. The Multi-faceted Ecto-enzyme CD38: Roles in Immunomodulation, Cancer, Aging, and Metabolic Diseases. Front Immunol. 2019;10.

37. Wei W, Graeff R, Yue J. Roles and mechanisms of the CD38/cyclic adenosine diphosphate ribose/Ca(2+) signaling pathway. World J Biol Chem. 2014;5(1):58–67.

38. Okabe K, Yaku K, Tobe K, Nakagawa T. Implications of altered NAD metabolism in metabolic disorders. J Biomed Sci. 2019;26(1):34.

39. Minhas PS, Liu L, Moon PK, et al. Macrophage de novo NAD+ synthesis specifies immune function in aging and inflammation. Nat Immunol. 2019;20(1):50–63.

40. Kang BN, Tirumurugaan KG, Deshpande DA, et al. Transcriptional regulation of CD38 expression by tumor necrosis factor-alpha in human airway smooth muscle cells: role of NF-kappaB and sensitivity to glucocorticoids. FASEB J. 2006;20(7):1000–2.

41. Covarrubias AJ, Kale A, Perrone R, et al. Senescent cells promote tissue NAD(+) decline during ageing via the activation of CD38(+) macrophages. Nat Metab. 2020;2(11):1265–83.

42. Chini CCS, Peclat TR, Warner GM, et al. CD38 ecto-enzyme in immune cells is induced during aging and regulates NAD(+) and NMN levels. Nat Metab. 2020;2(11):1284–304.

43. Shi B, Wang W, Korman B, et al. Targeting CD38-dependent NAD+ metabolism to mitigate multiple organ fibrosis. iScience. 2021;24(1).

44. Tao Y, Wang J, Lyu X, et al. Comprehensive Proteomics Analysis Identifies CD38-Mediated NAD(+) Decline Orchestrating Renal Fibrosis in Pediatric Patients With Obstructive Nephropathy. Mol Cell Proteomics. 2023;22(3):100510.

45. Tarragó MG, Chini CCS, Kanamori KS, et al. A Potent and Specific CD38 Inhibitor Ameliorates Age-Related Metabolic Dysfunction by Reversing Tissue NAD+ Decline. Cell Metab. 2018;27(5):1081–95.e10.

46. Becherer JD, Boros EE, Carpenter TY, et al. Discovery of 4-Amino-8-quinoline Carboxamides as Novel, Submicromolar Inhibitors of NAD-Hydrolyzing Enzyme CD38. J Med Chem. 2015;58(17):7021–56.

47. Barraco D, Cerquozzi S, Gangat N, et al. Monocytosis in polycythemia vera: Clinical and molecular correlates. Am J Hematol. 2017;92(7):640–5.

48. Tefferi A, Shah S, Mudireddy M, et al. Monocytosis is a powerful and independent predictor of inferior survival in primary myelofibrosis. Br J Haematol. 2018;183(5):835–8.

49. Fisher DAC, Malkova O, Engle EK, et al. Mass cytometry analysis reveals hyperactive NF Kappa B signaling in myelofibrosis and secondary acute myeloid leukemia. Leukemia. 2017;31(9):1962–74.

50. Laranjeira ABA, Kong T, Snyder SC, et al. In vivo ablation of NFκB cascade effectors alleviates disease burden in myeloproliferative neoplasms. Blood. 2024.

51. Matalonga J, Glaría E, Bresque M, et al. The Nuclear Receptor LXR Limits Bacterial Infection of Host Macrophages through a Mechanism that Impacts Cellular NAD Metabolism. Cell Reports. 2017;18:1241–55.

52. Cui H, Xie N, Banerjee S, et al. CD38 Mediates Lung Fibrosis by Promoting Alveolar Epithelial Cell Aging. Am J Respir Crit Care Med. 2022;206(4):459–75.

53. Schneider RK, Mullally A, Dugourd A, et al. Gli1(+) Mesenchymal Stromal Cells Are a Key Driver of Bone Marrow Fibrosis and an Important Cellular Therapeutic Target. Cell Stem Cell. 2017;20(6):785–800.e8.

54. Decker M, Martinez-Morentin L, Wang G, et al. Leptin-receptor-expressing bone marrow stromal cells are myofibroblasts in primary myelofibrosis. Nat Cell Biol. 2017;19(6):677–88.

