## supplementary file for "Targeting CD38 effectively prevents myelofibrosis in myeloproliferative neoplasms"

**Supplementary Materials for “Targeting CD38 effectively prevents myelofibrosis in myeloproliferative neoplasms”**

**Supplementary Methods**

**Mice**

For transplantation assay, 8-week-old CD45.1^+^ BL6.SJL recipient mice were lethally irradiated(4.5Gy×2) and transplanted with 3x10^6^ bone marrow nucleated cells(BMNCs) intravenously in 100μL PBS from *Mx1*-Cre (WT), *Nras^G12D/+^Mx1*-Cre (N), *Jak2^V617F/+^Mx1*-Cre (J) and *Nras^G12D/+^Jak2^V617F/+^Mx1*-Cre (NJ) CD45.2^+^ donor mice after poly(I:C) induction for 4-6 weeks. Whole blood cell counts were measured every 4 weeks until 28 weeks after transplantation.

For *in vivo* drug studies, 8-week-old CD45.1^+^ BL6.SJL recipient mice were lethally irradiated(4.5Gy×2) and transplanted with 3x10^6^ BMNCs from NJ CD45.2^+^ donor mice and randomized into treatment groups 20 weeks post-transplantation according to blood counts and body weight. Mice were treated with control (solvent) and 78c (30 mg/kg/d) by oral gavage for 4 weeks. The dosage of 78c was selected according to previous research^1, 2^.

Mice were housed in the animal barrier facility at the State Key Laboratory of Experimental Hematology. The experimental protocols were approved by the Institutional Animal Care and Use Committee of State Key Laboratory of Experimental Hematology.

**Blood and tissue histological analyses**

Peripheral blood (PB) was collected from the retro-orbital vein and measured by a Blood Cell Analyzer (Sysmex, XN-1000). Mouse tissue samples were fixed in 10% neutral-buffered formalin for 24 h and embedded in paraffin. Paraffin blocks were sectioned at 5 μm and stained with hematoxylin and eosin (H&E), reticulin staining according to the manufacturer’s instructions.

**Colony assays**

BMNCs (20,000 cells per well) or spleen nucleated cells (SPLNCs) (100,000 cells per well) were plated in triplicate using semisolid medium (MethoCult M3434, Stem Cell Technologies). On the seventh day, the number of colonies formed by burst-forming unit-erythroid (BFU-E), granulocyte-macrophage colony forming unit (CFU-GM), and colony-forming unit-granulocyte, erythrocyte, macrophage and megakaryocyte (CFU-GEMM) were counted.

***In vitro* fibrocyte differentiation assay and quantification**

Murine BMNCs or patients’ PB CD14^+^ monocyte were resuspended in conditions that promote the differentiation of monocytes to fibrocytes^3, 4^. Cells were cultured in flat-bottomed 24-well tissue culture plates with 5×10^5^

cells/500μL in a humidified incubator containing 5% CO_2_ at 37°C. BMNCs were first co-cultured with 78c or NMN for 24 hours. After 5 days, immunofluorescence staining was performed to identify fibrocytes as previously described^3^.

Representative microscopy images were captured using confocal microscopy (PerkinElmer UltraVIEW VoX system) with a magnification of 20×.

For the quantitative analysis of fibrocytes, 13 corresponding fields were captured by high content analysis systems (PerkinElmer) with a magnification of 20× for each well to measure fibrocytes (mouse:CD45^+^CollagenI^+^DAPI^+^; human: CD45^+^ Procollagen I^+^ DAPI^+^) counts using associated software.

**Treatment of fibrocytes with reagents and drugs *in vitro***

Murine cultured BM–derived fibrocytes were initially cultured in Roswell Park Memorial Institute (RPMI) 1640 medium supplemented with 10% fetal bovine serum (FBS), and exposed to 78c(MCE), NMN (sigma) for the first 24 hours. Following this, the culture medium was replaced with drug-free fibrocyte medium, and cells were cultured for 5 days. 78c were dissolved in DMSO (stock solution of 10 mM) and further diluted in the culture medium.

**Flow cytometry**

BMNCs from murine femurs and tibias were isolated by flushing them into PBS containing 2% Fetal Bovine serum (FBS, Gibco) and 2 mM EDTA (Invitrogen). Spleen cells were prepared by crushing and passing the tissue through a 70 μm cell strainer (Biosharp). PB was collected from the retro-orbital plexus. Single-cell suspensions of BM, spleen and PB were incubated for 30 min at 4°C with antibody cocktail in a total volume of 100μL PBS/2% FBS.

For surface staining, the following antibodies were used: CD45.1 (A20), CD45.2 (104), Lineage Antibody Cocktail (17A2, RB6-8C5, RA3-6B2, Ter-119, M1/70), CD117 (c-kit, 2B8), Ly-6A/E (Sca-1, D7), CD34 (RAM34), CD16/CD32 (FcγRII/III, 93), CD48(HM48-1), CD150(TC15-12F12.2), CD45R/B220 (RA3-6B2), CD3 (17A2), Ly6C(AL-21), Ly6G (Gr-1, RB6-8C5), CD11b (M1/70), F4/80 (BM8), CD115 (AFS98), CD80 (16-10A1), CD206 (C068C2), CD71(RI7217), TER-119 (TER-119), I-A/I-E (MHC class II, M5/114.15.2), CD11c(N418), CD4(GK1.5), CD135(A2F10), CD41(MWReg30), CD61(2C9.G2) and CD38(90). DAPI (Beyotime) or 7-AAD (BD Biosciences) were added to exclude dead cells before measurement. All FACS antibodies were purchased from BD Biosciences or Biolegend.

For intracellular collagen staining, the following antibodies were used: CD68 (FA-11), Collagen Type I Antibody Biotin Conjugated (Rockland), APC streptavidin. Cells were fixed and permeabilized using the IntraSure Kit (BD Biosciences). Viability was tracked using the Fixable Viability Dye eFluor 506 (Invitrogen, 65-0866-14) according to the manufacturer’s protocol.

All analyses were performed on a FACS Canto II flow cytometer (BD Biosciences) and analyzed with FlowJo software V10.

**Western blot**

Murine BMNCs were lysed with 2x Laemmli Sample Buffer (Bio-Rad), and total protein separated on PAGE gels (GenScript ProBio). Blots were probed for RAS G12D (Cell Signaling Technology, 14429) and Gapdh (BOSTER, BM1623).

**TNF-α enzyme-linked immunosorbent assay**

Serum TNF-α level in mice were measured using Mouse TNF-α ELISA Kit (EM0183, FineTest) following the manufacturer's protocol.

**NAD assay**

NAD^+^ level was measured by using the NAD^+^/NADH ratio Assay Kit (S0175, S0176S, Beyotime) and BCA Protein Assay Kit (BL521S, Biosharp) according to the manufacturer instructions

**RNA extraction and real-time quantitative PCR analysis**

Murine monocytes were sorted using sorted using FACS Aria III flow cytometer (BD Biosciences). Murine BM CD115^+^ cells were isolated and enriched using PE MicroBeads (Miltenyi) and PE anti-mouse CD115 (CSF-1R) Antibody (Biolegend) and separated using MACS separation column (Miltenyi). Total RNA from murine BM cells or human BMMNCs was purified using TRIzol (Invitrogen) and cDNA was reverse transcribed from 1μg total RNA using RevertAid First Strand cDNA Synthesis Kit (Thermo Fisher Scientific). Real-time quantitative PCR (RT-qPCR) was performed using the PowerUpTM SYBRTM Green Master Mix (Applied Biosystems) on a StepOne Real-Time PCR System (Thermo Fisher Scientific) and analyzed with associated software. Gapdh/GAPDH was used to normalize the RNA content of samples. Some of the primers were obtained from Primer Bank listed in Supplementary Table1 (https://pga.mgh.harvard.edu/primerbank/). All PCRs were conducted in triplicate.

**RNA sequencing and bioinformatics**

RNA amplification, library production, and data preprocessing were conducted by Novogene Co (Beijing, China). Briefly, a total amount of 1 μg RNA per sample was used for the RNA sample preparations. RNA libraries were generated using the NEBNext® UltraTM RNA Library Prep Kit for Illumina (NEB) following the manufacturer’s recommendations, and index codes were added to attribute sequences to each sample. All RNA-seq libraries were sequenced on an Illumina Novaseq platform and 150 bp paired-end reads were generated.

For RNA-seq analyses, sequence reads were aligned to the reference genome using Hisat2 v.2.0.5 after removing reads containing adapter and ploy-N, and low-quality reads. FeatureCounts v.1.5.0-p3 was used to count the reads mapped to each gene. Next, differential expression analysis of the two groups was conducted using the DESeq2 R package (1.16.1). Differentially expressed genes were selected based on a fold change >1 and *P* < 0.05.

**Quasi-Targeted Metabolomics**

Murine monocytes were sorted using sorted using FACS Aria III flow cytometer (BD Biosciences). Cells were washed once with pre-cooled PBS, followed by centrifugation at 1000g for 1 minute. The supernatant was carefully removed, and the cell pellet was snap-frozen in liquid nitrogen for 15 minutes. Subsequent metabolite analysis was performed by Novogene Co (Beijing, China).

**Statistical analysis**

Statistical analyses were done using GraphPad Prism software. All data are shown as mean ± standard error of the mean. The statistical significance of the two groups in the *in vivo* and *in vitro* trials was determined using an unpaired Student's t-test. Mann-Whitney U test was used for mRNA expression analysis of individuals. Kaplan-Meier survival analysis and log-rank test were used to compare survivals. Significance was set as **P* ≤ .05; ***P* ≤ .01; ****P* ≤ .001; *****P* ≤ .0001.

**References**

1.	Qiu Y, Xu S, Chen X, et al. NAD(+) exhaustion by CD38 upregulation contributes to blood pressure elevation and vascular damage in hypertension. Signal Transduct Target Ther. 2023;8(1):353.

2.	Shi B, Wang W, Korman B, et al. Targeting CD38-dependent NAD+ metabolism to mitigate multiple organ fibrosis. iScience. 2021;24(1).

3.	Shi Z, Liu J, Zhao Y, et al. ASXL1 mutations accelerate bone marrow fibrosis via EGR1-TNFA axis mediated neoplastic fibrocyte generation in myeloproliferative neoplasms. Haematologica. 2022.

4.	Ozono Y, Shide K, Kameda T, et al. Neoplastic fibrocytes play an essential role in bone marrow fibrosis in Jak2V617F-induced primary myelofibrosis mice. Leukemia. 2021;35(2):454-67.

**Supplemental Table 1. qRT-PCR Primers**

**For mouse**

| **Genes** | **Sequence** |
| --- | --- |
| Gapdh-F | CGTCCCGTAGACAAAATGGT |
| Gapdh-R | TTGATGGCAACAATCTCCAC |
| Cd38-F | CATCACAAGAGAAGACTACGCCCC |
| Cd38-R | CACACCACCTGAGATCATCAGCAA |
| Cd157-F | TGCTCGTTATGAGCTATGGGG |
| Cd157-R | TCAAGTCCAGAGGCATTTTCC |
| Sirt1-F | GAACCACCAAAGCGGAAAAAAAG |
| Sirt1-R | AATCCCACAGGAGACAGAAACCC |
| Sirt3-F | CTCAAAGCTGGTTGAAGCCCACG |
| Sirt3-R | GCTCCCCAAAGAACACAATGTCG |
| Parp1-F | AAGAAAGGGAAGGACAAGGATAG |
| Parp1-R | CACCTGCTGCTGGTTGAAGATGA |
| Parp2-F | GCAACAGAAGACGACTCTCCT |
| Parp2-R | CAGCCATAGGCCCTTTTCTCT |
| Nampt-F | CATAGGGGCATCTGCTCATTTGGTT |
| Nampt-R | ATGGTACTGTGCTCTGCTGCTGGAA |
| Nmnat1-F | TGGCTCTTTTAACCCCATCAC |
| Nmnat1-R | TCTTCTTGTACGCATCACCGA |
| Nmnat3-F | TCACCCGTCAATGACAGCTAT |
| Nmnat3-R | CACCCGAATCCAGTCAGATGT |
| Ido1-F | GCCTCCTATTCTGTCTTATGCAG |
| Ido1-R | ATACAGTGGGGATTGCTTTGATT |

**For human**

| **Genes** | **Sequence** |
| --- | --- |
| GAPDH-F | CATGAGAAGTATGACAACAGCCT |
| GAPDH-R | AGTCCTTCCACGATACCAAAGT |
| CD38-F | TCCTGGTCCTGATCCTCGTCGT |
| CD38-R | CCCTTGAAAGCATCCCATACAC |
| CD157-F | ACTTGCGGGACATCTTCCTG |
| CD157-R | AAAGGTCATAGTCTGAGGGCA |
| SIRT1-F | TAGCCTTGTCAGATAAGGAAGGA |
| SIRT1-R | ACAGCTTCACAGTCAACTTTGT |
| SIRT3-F | ACCCAGTGGCATTCCAGAC |
| SIRT3-R | GGCTTGGGGTTGTGAAAGAAG |
| PARP1-F | CGGAGTCTTCGGATAAGCTCT |
| PARP1-R | TTTCCATCAAACATGGGCGAC |
| PARP2-F | GCCTTGCTGTTAAAGGGCAAA |
| PARP2-R | TCCTTCACAATACACATGAGCC |
| NAMPT-F | CGGCAGAAGCCGAGTTCAA |
| NAMPT-R | GCTTGTGTTGGGTGGATATTGTT |
| NMNAT1-F | TCTCCTTGCTTGTGGTTCATTC |
| NMNAT1-R | TGACAACTGTGTACCTTCCTGTT |
| NMNAT3-F | GAGTAGGTCACGACCCAAAAG |
| NMNAT3-R | TCGCCTGATGTATGTGGCAC |
| IDO1-F | GCCAGCTTCGAGAAAGAGTTG |
| IDO1-R | ATCCCAGAACTAGACGTGCAA |

**Supplemental Table 2.**

**Patients’ Laboratory Characteristics in the Validation of CD38 Expression Levels: Comparative Analysis of MPN Patients with and without Fibrosis Development during Follow-up (related to Figure 4l). (Last Follow-Up Date: 2024-08-31)**

| Patients# | Diagnosis | First Diagnosis Date | Fibrosis Development (Date) | Mutation (allele burden) |
| --- | --- | --- | --- | --- |
| **5257 | PV | 2018-07-18 | No | JAK2 p.V617F(75.6%), NOTCH1 p. Q1134R (47.5%) |
| **6975 | PV | 2018-05-24 | No | JAK2 p.V617F(35.3%), TET2 p.R1359H (27.2%) |
| **1713 | PV | 2018-06-25 | No | JAK2 p.V617F(47.2%),  KMT2D p. T2524M (54.5%), FAT1 p.L2822P (40.6%) |
| **7992 | PV | 2018-10-10 | No | JAK2 p.V617F(44.7%) |
| **4282 | PV | 2018-11-21 | No | JAK2 p.V617F(44.9%), CREBBP p.A254T (50.2%),  ARID1A p.P1771S (46.7%), MYD88 p.R230C (49.7%) |
| **5174 | PV | 2018-11-27 | No | JAK2 p.V617F (43.3%), KMT2D p. P1191L (47.3%),  PLCG1 p.I1092V (46.9%), FGFR3 p.L164V (48.5%) |
| **4042 | PV | 2019-04-08 | No | JAK2 p.V617F 61.1% |
| **6659 | PV | 2019-04-23 | No | JAK2 p.V617F (29.9%), FAT1 p.I3726N (52.3%) |
| **7887 | PV | 2019-07-01 | No | JAK2 p.V617F (33.4%), JAK2 p.G127D (32.9%) |
| **8259 | PV | 2019-07-05 | No | JAK2 p.V617F (53.5%),  SF3B1 p.K666N (24.5%), TET2 p.K1243R (48.2%) |
| **6047 | **PV** | 2018-05-21 | **Yes (2021-05-24)** | **2018-05-21(PV stage)**  JAK2 p.V617F(1.1%), JAK2 p.K539L(30.8%),  SETD2 p.R433C(49.0%), BRINP3 p.R6S(52.0%)  RELN p.M2185L(51.7%), KMT2D p.T2228A(45.4%) |
|  |  |  |  | **2021-05-24(post-PV MF stage)**  JAK2 p.K539* (47.0%), DNMT3A p.G543S (43.5%) |
| **9267 | **PV** | 2019-05-10 | **Yes (2022-09-19)** | **2019-05-10(PV stage)**  JAK2 p.V617F(40.1%), EZH2 p.H689R(7.6%), USP7 p.G392D(7.8%)  ASXL1 p.S1429del(49.7%), KMT2D p.S2215T(48.8%), TET2 p.F1300C(1.2%), ASXL1 p.P229L(49.5%) |
|  |  |  |  | **2022-09-19(post-PV MF stage)**  JAK2 p.V617F(43.0%), ASXL1 p.G646Wfs*12(27.8%), EZH2 p.R684C(30.9%), IKZF1 p.V341Sfs*74(3.8%), EZH2 p.H694R(31.7%) |
| **2014 | ET | 2014-05-21 | No | CALR p.K385Nfs*47(34.2%) |
| **1485 | ET | 2015-12-10 | No | JAK2 p.V617F(41.43%), TET2 p.H1904R(30.86%) |
| **9110 | ET | 2015-11-19 | No | SF3B1 p.L747W(1.89%), MPL p.R321Q(47.59%) |
| **3063 | ET | 2015-12-24 | No | JAK2 p.V617F(10.92%), NPM1 p.960dupTCTG |
| **6684 | **ET** | 2017-05-23 | **Yes (2021-11-28)** | **2017-05-23(ET stage)**  JAK2 p.V617F(88.0%), TET2 p.I1873T (43.3%), ATM p.Q1982R (48.6%), CREBBP p.A254T(48.1%), TET2 p.Y1345_N1346del (43.4%), FAT1 p.L2822P(51.5%) |
|  |  |  |  | **2021-11-28(post-ET MF stage)**  JAK2 p.V617F(96.3%), TET2 p.I1873T(47.7%), TP53 p.C135W(11.5%) |

**Supplementary Figure Legends**
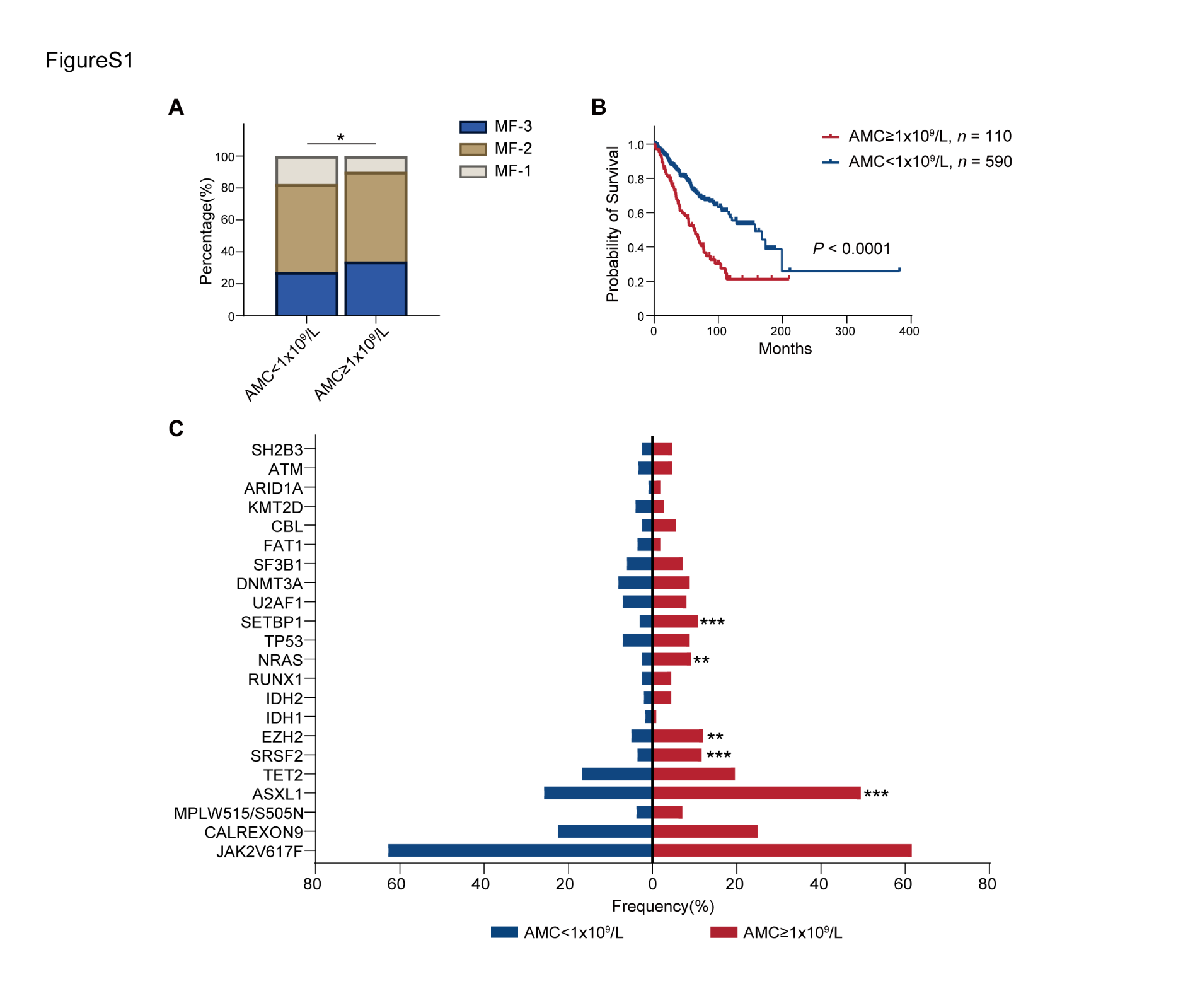

**Supplementary Figure 1**

**Figure S1. PMF patients with monocytosis are associated with severe disease phenotypes.**

**A.** Bone marrow fibrosis grades of MF patients with different absolute monocyte counts. (*n* = 603 for AMC<1x10^9^/L patients, *n* = 112 for AMC≥1x10^9^/L patients). **B.** Comparison of survival between MF patients with or without monocytosis (AMC≥1x10^9^/L). **C.** Genetic distribution of MF patients stratified by monocyte status. χ^2^ test or Fisher’s exact test were performed between percentages of 2 groups. AMC: absolute monocyte count. **P* ≤ .05; ***P* ≤ .01; ****P* ≤ .001; *****P* ≤ .0001.

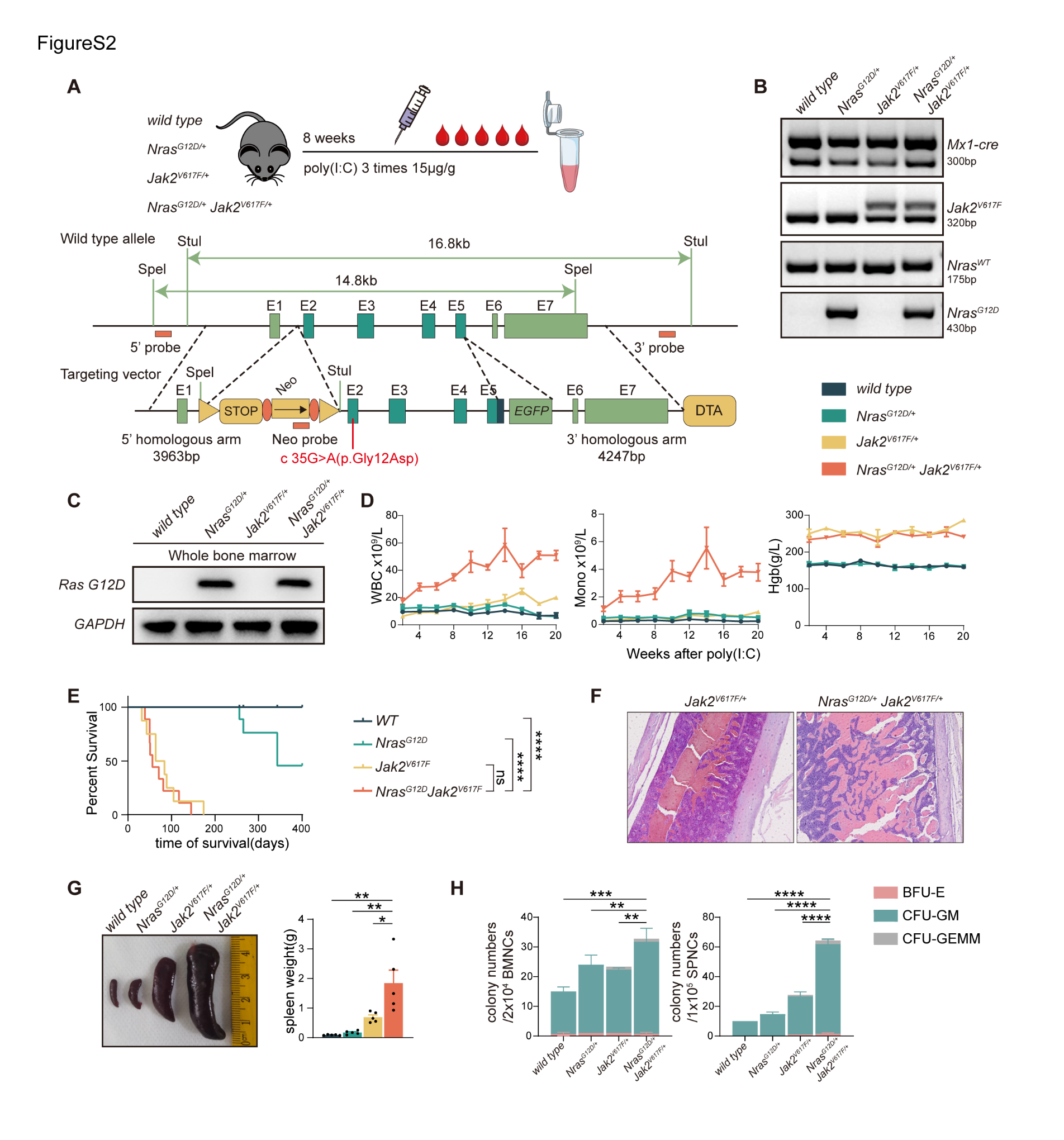
**Supplementary Figure 2**

**Figure S2. Phenotype of primary mouse.**

**A.** Schematic of the primary *Nras^G12D/+^Jak2^V617F/+^* -Mx1-Cre mouse model and targeting vector schema for *Nras^G12D/+^* mouse. **B.** PCR analysis of tail DNA from wild type, *Nras^G12D/+^*, *Jak2^V617F/+^*, and *Nras^G12D/+^Jak2^V617F/+^* mice. **C.** Verification of Ras G12D protein expression by western blot in whole bone marrow cells from WT, N, J, and NJ mice after 6 weeks of poly(I:C) treatment, with Gapdh as a loading control. **D.** White blood cell (WBC) counts, monocyte counts and hemoglobin level in wild type, *Nras^G12D/+^*, *Jak2^V617F/+^*, and *Nras^G12D/+^Jak2^V617F/+^* mice after poly(I:C). **E.** Survival analysis of wild type (*n* = 9), *Nras^G12D/+^* (*n* = 9), *Jak2^V617F/+^* (*n* = 8), and *Nras^G12D/+^Jak2^V617F/+^* (*n* = 9) mice. **F.** Hematoxylin and eosin (H&E) staining of bone marrow (BM) from representative *Jak2^V617F/+^* and *Nras^G12D/+^ Jak2^V617F/+^* mice demonstrated thrombosis in femur. Scale = 20 μm; Original magnification, 10×. The arrows indicate thrombus formation. **G.** Spleen size in the four groups of mice after 6 weeks of poly(I:C) treatment. **H.** Colony formation after 8 days of plating 20,000 BMNCs and 100,000 SPLNCs from wild type, *Nras^G12D/+^*, *Jak2^V617F/+^* and *Nras^G12D/+^ Jak2^V617F/+^* mice in methylcellulose containing rmIL-3, rm-SCF, rh-IL6, and rh-EPO (MethoCult M3434), with *n* = 3 per group. All data represent mean ± standard error of the mean (SEM). Statistics were assessed by two-tailed Student’s T test. **P* ≤ .05; ***P* ≤ .01; ****P* ≤ .001; *****P* ≤ .0001.

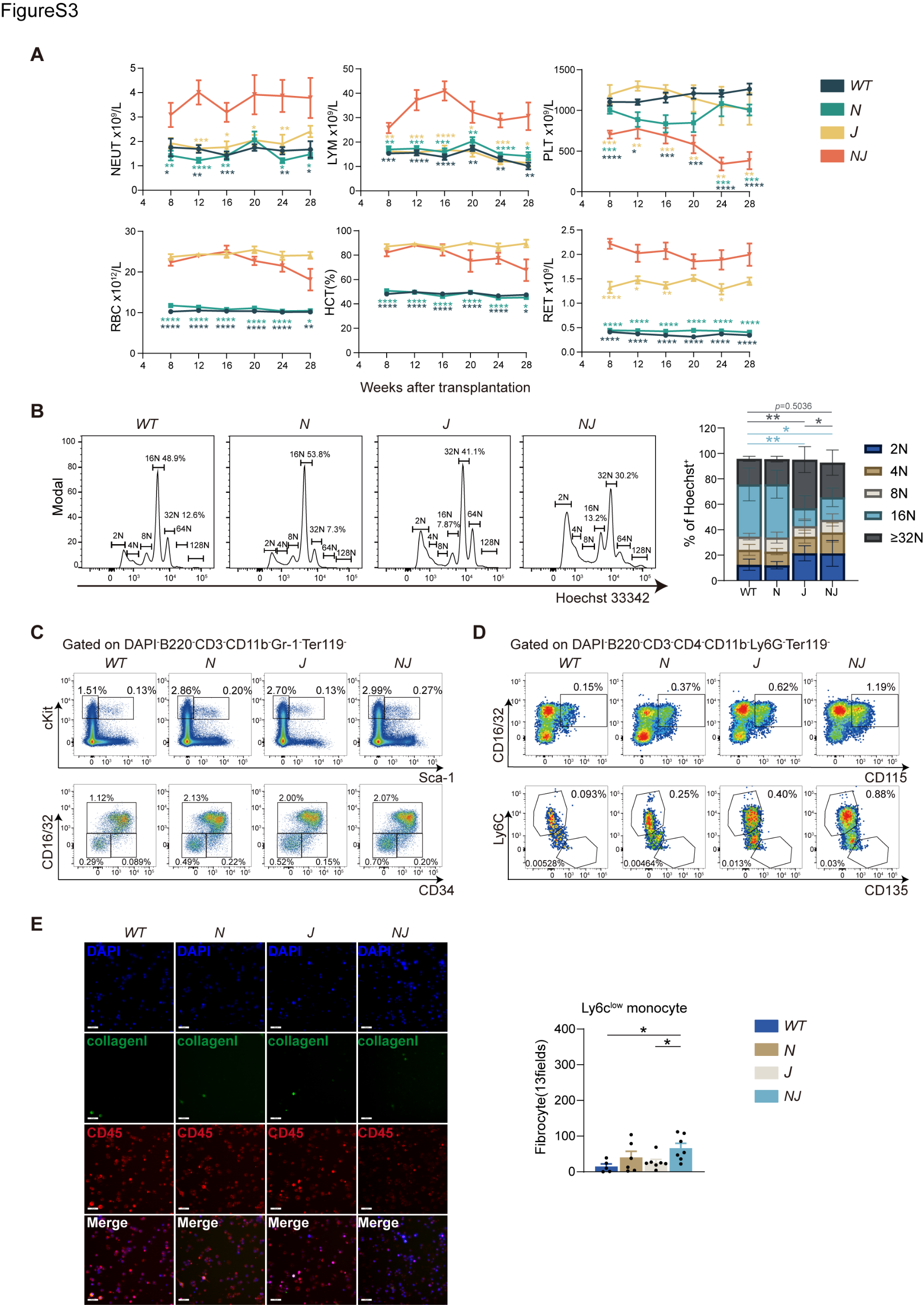
**Supplementary Figure 3**

**Figure S3. Phenotype in transplanted mouse.**

**A.** Neutrophil, lymphocyte, platelet, red blood cell (RBC) and reticulocyte (RET) counts and hematocrit (%) of wild type (WT, *n* = 10), *Nras^G12D/+^*(N, *n* = 8), *Jak2^V617F/+^*(J, *n* = 8), and *Nras^G12D/+^Jak2^V617F/+^* (NJ, *n* = 10) mice over 28 weeks post-transplantation. **B.** Ploidy analysis of megakaryocytes in bone marrow from WT, N, J and NJ mice 20 weeks post-transplantation. Gray asterisks represent the statistical comparison of ≥32-ploidy cells among the four groups and blue asterisks represent the statistical comparison of 16-ploidy cells among the four groups. **C.** Representative flow cytometric plots of Lin^−^Sca-1^+^c-Kit^+^ (LSK) and Lin^−^ckit^+^Sca-1^−^ (myeloid progenitor, MP), common myeloid progenitor (CMP, Lin^−^Sca-1^−^c-kit^+^CD34^+^FcγRII/III^low^), granulocyte-monocyte progenitor (GMP, Lin^−^Sca-1^−^c-kit^+^CD34^+^FcγRII/III^high^). **D.** Representative flow cytometric plots of monocyte-dendritic cell progenitor (MDP, Lin^−^Sca-1^−^c-kit^+^CD115^+^FcγRII/III^high^ CD135^+^ Ly6c^-^), and common monocyte progenitor (cMOP, Lin^−^Sca-1^−^c-kit^+^CD115^+^FcγRII/III^high^CD135^-^Ly6c^+^ in WT, N, J, and NJ mice 20 weeks post-transplantation. **E.** Representative immunofluorescence imaging and numbers of cultured monocytes derived fibrocytes derived from Ly6c^low^ monocyte among four groups. Scale = 40 μm; Original magnification 20×. All data represent mean ± standard error of the mean (SEM). Statistics were assessed by two-tailed Student’s T test. **P* ≤ .05; ***P* ≤ .01; ****P* ≤ .001; *****P* ≤ .0001.

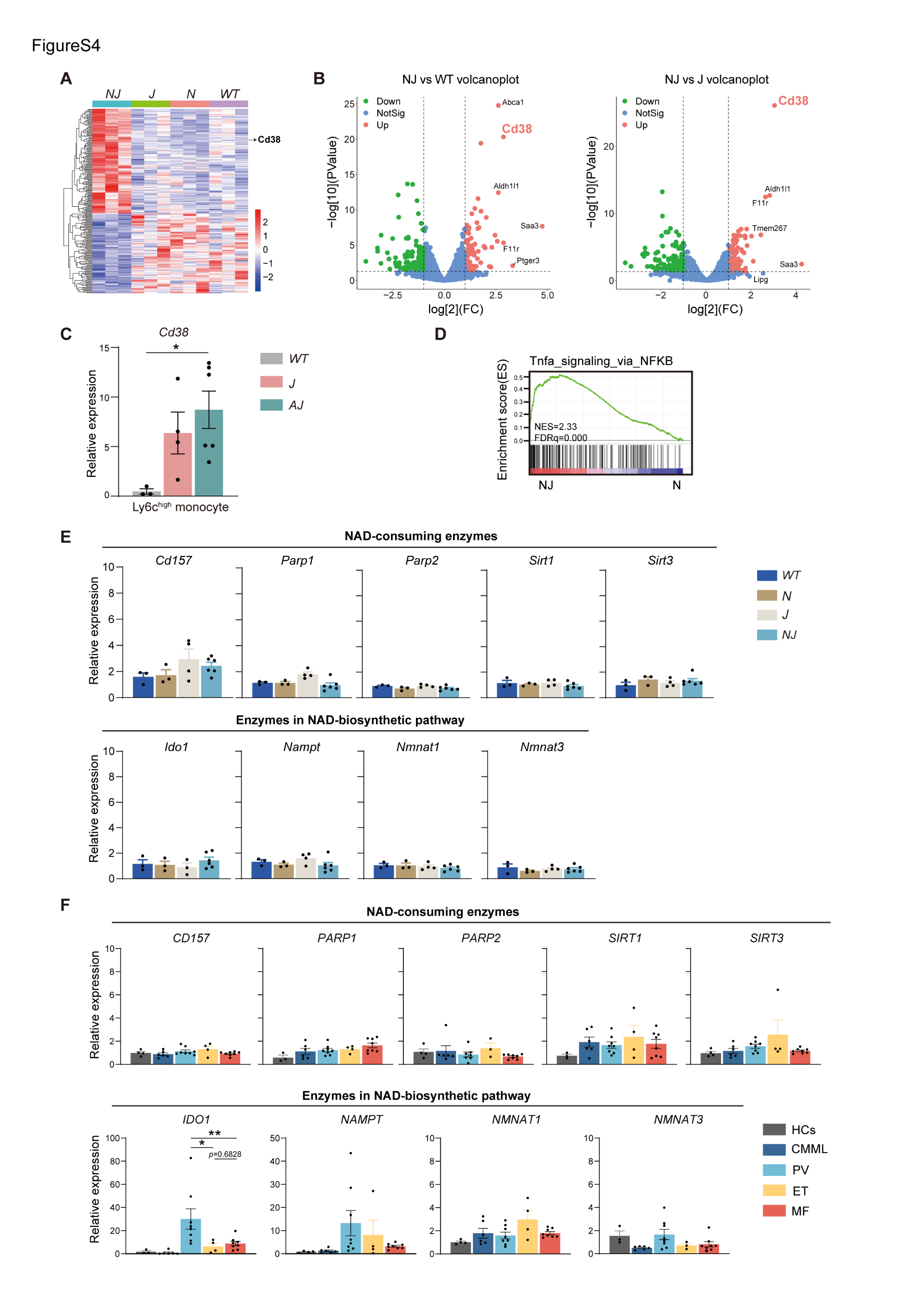
**Supplementary Figure 4**

**Figure S4. RNA-seq revealed *Cd38* overexpressed monocytes in NJ mice.**

**A.** Heat map of differentially expressed genes (DEGs) in NJ compared with J, N and WT BM CD11b^+^CD115^+^Ly6c^high^ monocyte (fold change≥1 fold; *p*≤ .05). **B.** Volcano plot(left) showing DEGs between NJ and WT. Volcano plot(right) showing DEGs between NJ and J. **C.** Quantification of mRNA levels of CD11b^+^CD115^+^Ly6c^high^ from *wild type* (WT), *Jak2^V617F^* Vav1-Cre (J), *Asxl1^-/-^ Jak2^V617F^* Vav1-Cre (AJ) mice at 30 weeks of age by qRT-PCR. **D.** Gene set enrichment analysis (GSEA) showed that enrichment gene sets between NJ and N. **E.** Quantification of mRNA levels of the enzymes involved in NAD^+^ biosynthesis or metabolism of in sorted BM CD11b^+^CD115^+^Ly6c^high^ monocyte from WT, N, J and NJ mice 20 weeks post-transplantation by qRT-PCR. **F.** Quantification of mRNA levels of the enzymes involved in NAD^+^ biosynthesis or metabolism of in human CD14^+^ monocytes by qRT-PCR. All data represent mean ± standard error of the mean (SEM). Statistics were assessed by two-tailed Student’s T test. **P* ≤ .05; ***P* ≤ .01; ****P* ≤ .001; *****P* ≤ .0001.

**Supplementary Figure 5**

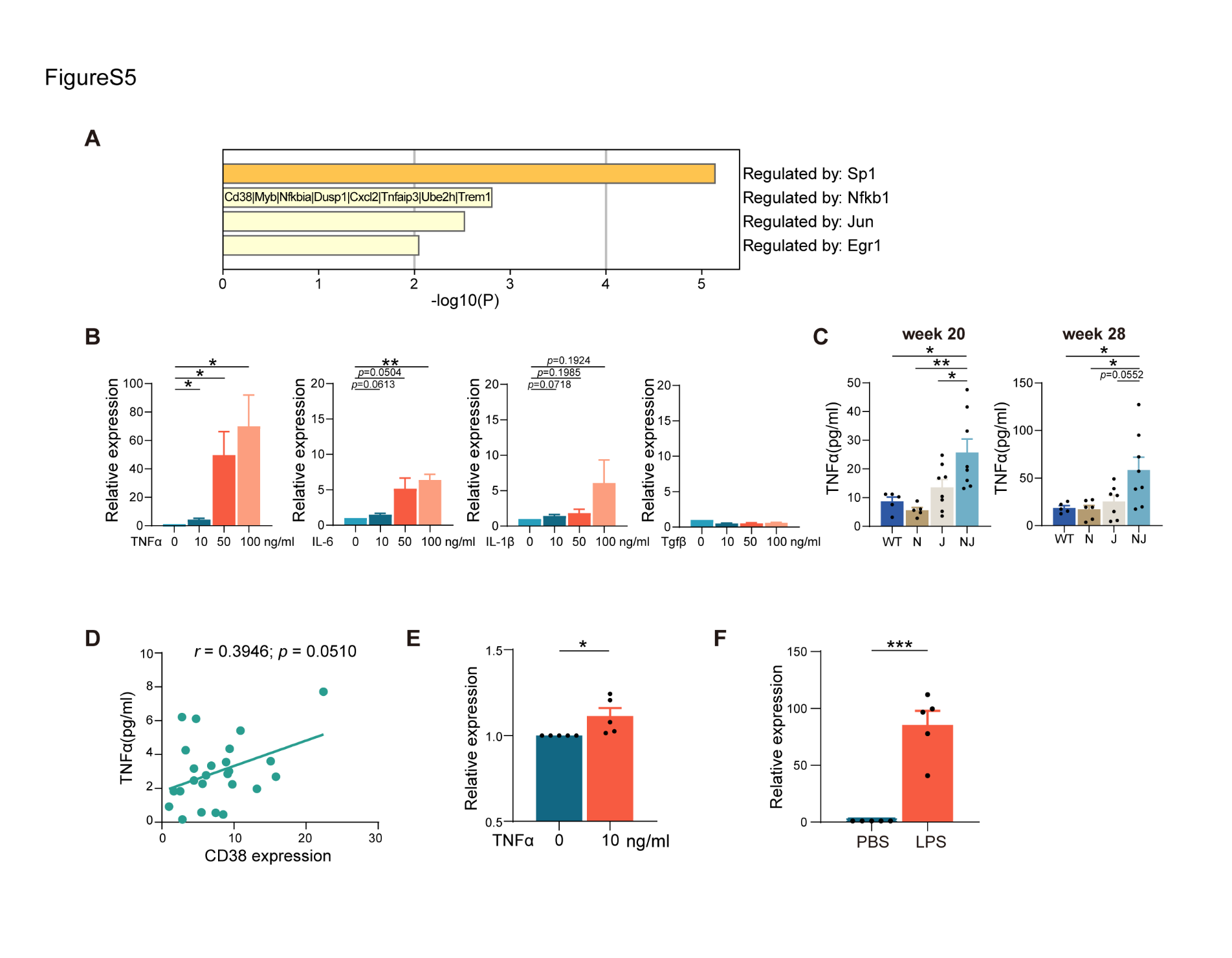

**Figure S5. TNF-α promotes increased CD38 expression in monocytes**

**A.** TRRUST (Transcriptional Regulatory Relationships Unraveled by Sentence-based Text mining) database analysis of 229 DEGs in NJ compared with J, N and WT BM CD11b^+^CD115^+^Ly6c^high^ monocyte (fold change ≥ 1 fold; *p* ≤ .05). **B.** Quantification of mRNA levels of *Cd38* in CD115^+^ monocytes following 16-hour exposure to TNF-α, IL-6, IL-1β and Tgfβ by qRT-PCR. **C.** Serum TNF-α level of four groups at 20 and 28 weeks. **D.** Correlation(*r*) between serum TNFα and CD38 expression level in MPN patients with *Jak2^V617F^* mutation (*n* = 25, PV *n* = 8; ET *n* = 4; MF *n* = 13). Pearson correlation noted. **E.** Patient (n = 6) monocytes were stimulated with human TNFα at the concentrations shown *in vitro*. After 24 hours of culture, monocyte was analyzed for QRT-PCR analysis. **F.** Quantification of the mRNA levels of *Cd38* in CD115^+^ monocytes from N mice after a 16-hour exposure to 100 ng/mL LPS by qRT-PCR. All data represent mean ± standard error of the mean (SEM). Statistics were assessed by two-tailed Student’s T test. **P* ≤ .05; ***P* ≤ .01; ****P* ≤ .001; *****P* ≤ .0001.

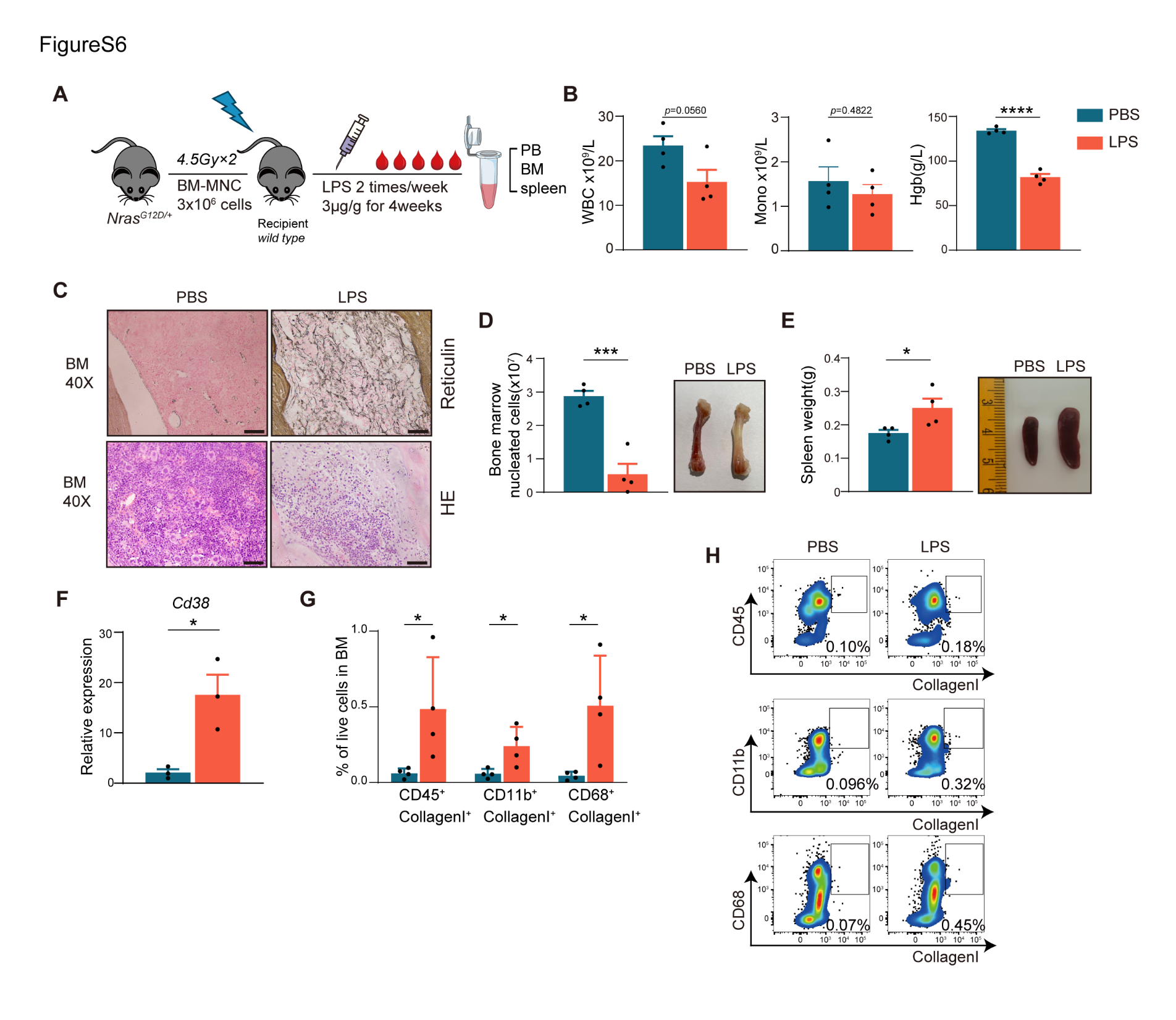
**Supplementary Figure 6**

**Figure S6. Overexpression of CD38 in monocyte induces bone marrow fibrosis**.

**A.** Schematic of LPS administration schedule in *Nras^G12D/+^* -Mx1-Cre transplanted mouse model induced. **B.** White blood cell (WBC) counts, monocyte counts and hemoglobin level of *Nras^G12D/+^* mice injected with PBS or LPS for 4 weeks. **C.** H&E and Reticulin staining of bone marrow (BM). Scale = 50 μm; Original magnification 40×. **D.** Bone marrow cell count per femur of *Nras^G12D/+^* mice injected with PBS or LPS for 4 weeks. **E.** Spleen size and weight of *Nras^G12D/+^* mice injected with PBS or LPS for 4 weeks. **F.** Quantification of mRNA levels of *Cd38* in sorted BM CD11b^+^CD115^+^Ly6c^high^ monocyte by qRT-PCR. **G.** The percentage of fibrocytes of two groups. **H.** The representative flow cytometry dot plot of fibrocytes of two groups. All data represent mean ± standard error of the mean (SEM). Statistics were assessed by two-tailed Student’s T test. **P* ≤ .05; ***P* ≤ .01; ****P* ≤ .001; *****P* ≤ .0001.

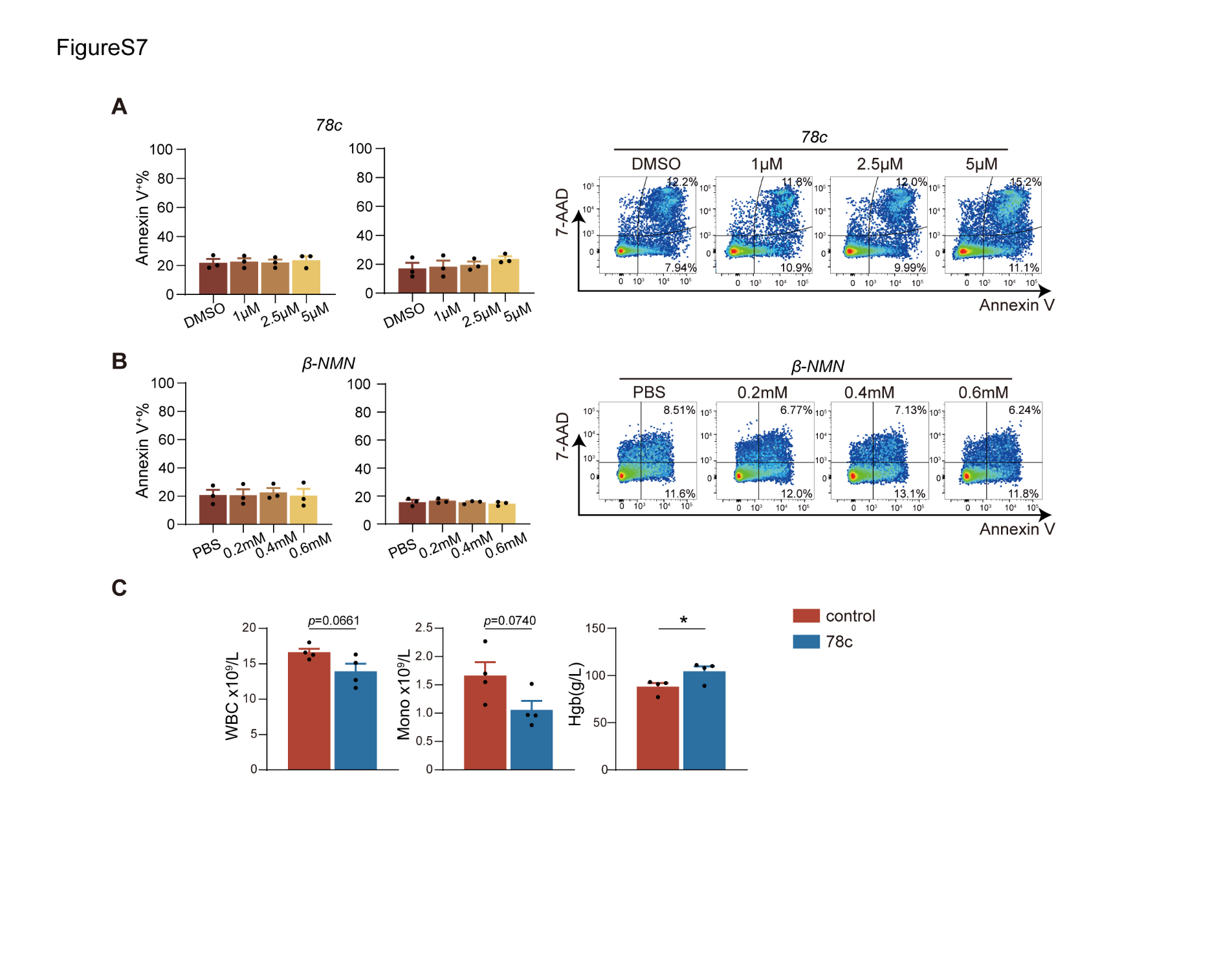
**Supplementary Figure 7**

**Figure S7. Effect of 78c and NMN on *Nras^G12D/+^Jak2^V617F/+^*** **bone marrow cells and hematological parameters of LPS-induced *Nras^G12D/+^* -Mx1-Cre model treated with either solvent or 78c.**

**A-B.** Quantification of apoptotic cell fractions in bone marrow cell cultures treated with 78c or NMN for 24 hours (left). Following this, the culture medium was replaced with drug-free medium, and cells were incubated for an additional 24 hours before apoptosis assessment (right). **C.** White blood cell (WBC) counts, monocyte counts and hemoglobin level of LPS-induced *Nras^G12D/+^* -Mx1-Cre transplanted mouse model treated with either solvent or 78c. All data represent mean ± standard error of the mean (SEM). Statistics were assessed by two-tailed Student’s T test. **P* ≤ .05; ***P* ≤ .01; ****P* ≤ .001; *****P* ≤ .0001.
